# Loss of metabolic resilience underlies maladaptive responses to lipid overload in TDP-43 proteinopathy

**DOI:** 10.64898/2026.11.15.766742

**Authors:** Juan M. Godoy-Corchuelo, Irene García-Toledo, Juan I. López-Carbonero, Angélica Larrad-Sainz, Zeinab Ali, María J. Gil-Moreno, Laura Fernández-Hernández, Luis C. Fernández-Beltrán, Lidia Peña-de-Diego, Irene Jiménez-Coca, Jesús Jiménez-Rodríguez, Rubén Fernández-de-la-Rosa, Ana Fernández-Bernal, Manuel Portero-Otin, María J. Casarejos-Fernández, Alicia Mansilla-Aparicio, Jimena Baleriola, Elizabeth M. C. Fisher, Jordi A. Matias-Guiu, Pablo Bascuñana, Thomas J. Cunningham, Silvia Corrochano

## Abstract

**INTRODUCTION:** Altered eating behaviour is a hallmark feature of behavioural variant frontotemporal dementia (bvFTD). These changes may represent a compensatory response to an underlying metabolic disturbance. However, whether increased lipid availability restores metabolic homeostasis or instead accelerates disease progression remains unknown. TDP-43 proteinopathies, which account for approximately half of FTD cases, are associated with alterations in lipid metabolism, providing a mechanistic framework to investigate the cerebral response to lipid overload.

**METHODS:** We combined clinical observations in patients with bvFTD with mechanistic studies in the physiological *Tardbp^M323K/M323K^*knock-in mouse model to determine how TDP-43 dysfunction modifies the response to chronic lipid excess.

**RESULTS:** In the bvFTD cohort, elevated cholesterol and triglycerides were associated with poorer cognitive performance, while mutant mice fed a lifelong high-fat diet showed worse cognitive impairment. Multi-omics analyses of the frontal cortex revealed maladaptive metabolic rewiring characterized by impaired mitochondrial and cholesterol metabolism, lipid accumulation, and widespread remodelling of TDP-43-dependent RNA processing. These alterations affected synaptic function and neuronal communication and were accompanied by reduced cerebral glucose utilization measured by ^18^F-FDG PET, neurotransmitter imbalance and enhanced neuroinflammation.

**DISCUSSION:** Increased lipid availability fails to compensate for TDP-43-associated metabolic dysfunction and instead promotes maladaptive molecular and functional responses. These findings identify impaired metabolic resilience as feature of TDP-43-mediated neurodegeneration and suggest that altered eating behaviour in bvFTD may represent an unsuccessful attempt to counteract an underlying metabolic deficit rather than a purely behavioural symptom. Thus, the biological context may determine whether increased energy intake is beneficial or detrimental during neurodegeneration.

## Background

Neurodegenerative diseases leading to dementia, including Alzheimer’s disease (AD) and frontotemporal dementia (FTD), represent a major and growing global health burden (1). As evidence indicates that dietary and other lifestyle factors influence dementia risk, nutrition-focused prevention strategies are increasingly being incorporated into large international initiatives such as Euro-FINGERS (2). However, most dietary interventions have been developed largely within an AD-centric framework and implicitly assume shared metabolic vulnerabilities across neurodegenerative disorders (3). Increasing evidence suggests instead that distinct molecular pathologies may differentially shape brain metabolism and modify responses to nutritional interventions. This highlights the need for etiology-specific approaches, particularly in TDP-43 proteinopathies, which may have metabolic and therapeutic requirements distinct from those of classical AD (4).

TDP-43 is a central pathological protein in a broad spectrum of neurodegenerative disorders. Loss of its nuclear functions and cytoplasmic aggregation lead to widespread transcriptomic dysregulation, impaired proteostasis, and metabolic vulnerability. TDP-43 pathology is present in approximately 97% of amyotrophic lateral sclerosis cases and around 50% of FTD cases, while also occurring as a co-pathology in several other neurodegenerative diseases, including AD (5), Parkinson’s disease, Huntington’s disease (6), and limbic-predominant age-related TDP-43 encephalopathy (LATE) (7). We and others have shown that TDP-43 dysfunction disrupts cholesterol and myelin biology (8), impairs lipid handling (8,9), and alters fatty-acid and mitochondrial metabolism (10). These metabolic disturbances are increasingly recognized as important contributors to neuronal and glial dysfunction across TDP-43 proteinopathies.

Diet is a major environmental determinant of neurodegenerative risk and a potentially modifiable factor capable of altering disease trajectory. In particular, Western-style diets enriched in saturated fats and ultra-processed foods have been associated with cognitive decline and dementia (11), although studies examining the effects of high-fat diets on brain function have yielded inconsistent results across experimental and clinical settings (12).

These relationships are particularly intriguing in frontotemporal dementia (FTD). In contrast to Alzheimer’s disease, where weight loss frequently precedes or accompanies disease progression, several studies have reported increased body mass index, weight gain, and broader metabolic alterations in patients with FTD (13). Once behavioural variant FTD (bvFTD) becomes clinically manifest, profound alterations in feeding behaviour are commonly observed, including hyperphagia, excessive caloric intake, a marked preference for sweet and energy-dense foods, often associated with weight gain (14). These features have traditionally been interpreted as a behavioural consequence of degeneration affecting hypothalamic, frontoinsular, and reward-related circuits that regulate feeding and energy homeostasis (15). However, mechanistic studies indicate that such alterations are particularly prominent with underlying TDP-43 pathology, raising the possibility that they may also reflect disease-specific metabolic adaptations (16). Indeed, growing evidence indicates that TDP-43 dysfunction disrupts cholesterol homeostasis, lipid handling, and cellular bioenergetics.

An intriguing possibility is that the changes in eating behaviour observed in bvFTD represented adaptive responses to an underlying lipid metabolic dysregulation. If so, increased lipid availability could help restore energetic balance and mitigate disease-associated dysfunction. Alternatively, these metabolic alterations may reflect a pre-existing disease-related vulnerability, and so increased lipid exposure could further challenge an already compromised metabolic system. Distinguishing between these possibilities is essential for the development of mechanism-based nutritional interventions, as dietary strategies that are beneficial in one neurodegenerative context may prove ineffective or even harmful in another. To address this knowledge gap, it is necessary to identify the pathways that mediate the effects of altered lipid metabolism on neuronal function and cognition, which will help determine the biological basis of metabolic susceptibility and guide the development of precision nutritional and therapeutic interventions.

Here, we combined clinical analyses in patients with bvFTD and experimental studies in the *Tardbp*-M323K knock-in mouse model, which recapitulates key features of frontotemporal dementia (17). Using this model, we have previously shown that TDP 43 dysfunction perturbs brain lipid metabolism (8). Here, we expose these mice to a lifelong high-fat dietary intervention together with comprehensive behavioural, neuroimaging, and multi-omics approaches to determine whether and how increased lipid availability compensates for, or exacerbates, the metabolic consequences of TDP-43 dysfunction.

## Results

### 1. Increased blood lipids were associated with worse cognitive outcomes in bvFTD patients

To investigate whether altered food intake and weight changes in behavioural variant frontotemporal dementia (bvFTD) could represent beneficial or detrimental adaptive responses to an underlying “metabolic vulnerability”, we evaluated the metabolic alterations and changes in the eating behaviour in a cross-sectional study of patients with bvFTD (n=30). Demographic data from these patients can be found in Table 1. First, we evaluated changes in their eating behaviour (hyperorality, changes in food preference, binge eating, and substance use/increased toxicity). The majority of the patients presented with altered eating behaviours (27 out of 30). Nevertheless, systemic metabolic changes were heterogeneous according to weight variations from the premorbid weight-5 years before symptom onset. Some patients gained weight (n=17) but also there were some who lost weight (n=13) (Table 1 and Fig.1a), and those differences were not associated with the sex (Table 1). The caloric intake did not differ significantly between those who gained or lost weight, inferred from a 24-hour food recall questionnaire (Fig. 1b). There were no clear differences in macronutrient intake (carbohydrates or lipids) or in the distribution of lipid species according to their degree of unsaturation (i.e., the number of double bonds in their acyl chains) (Fig. S1a,b).

**Figure 1.**
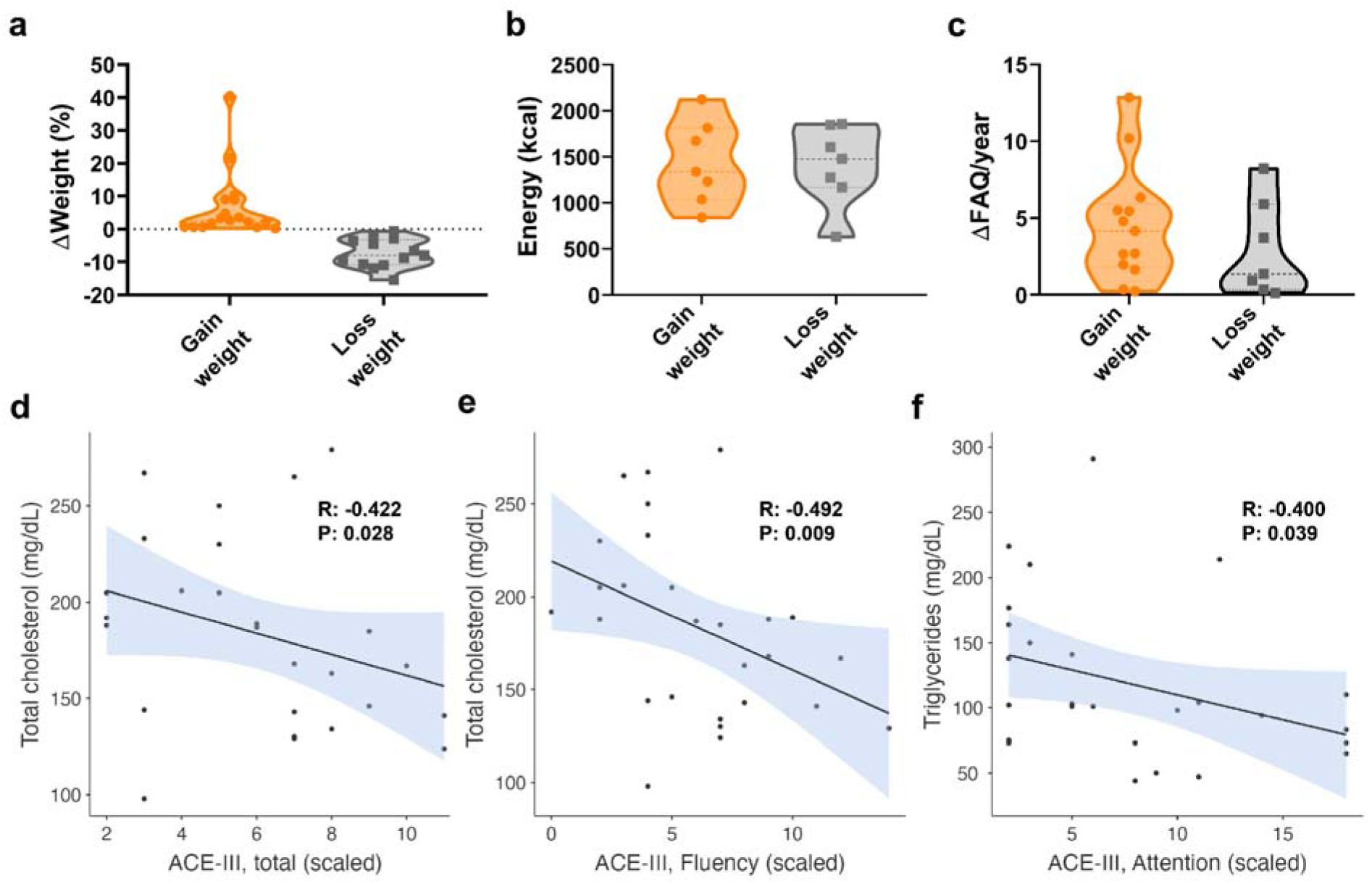
Metabolic and clinical characterization of FTD patients stratified by weight dynamics. **a.** Bar plot depicting the distribution of the FTD cohort categorized by weight trajectory: weight gain vs. weight loss. **b.** Comparative analysis of total daily caloric intake (kcal/day) between weight-gain and non-weight-gain cohorts. **c.** Longitudinal progression of functional decline, measured as the annual delta FAQ rate. **d**. Correlation between Total cholesterol levels and ACE-II total (scaled) score (Rho= -0.422 and p value= 0.028). **e**. Correlation between Total cholesterol levels and ACE-III, Fluency (scaled) score (Rho= -0.492 and p value= 0.009). **f**. Correlation between Triglycerides levels and ACE-III, Attention (scaled) score (Rho= -0.400 and p value= 0.039).

**Table 1.**
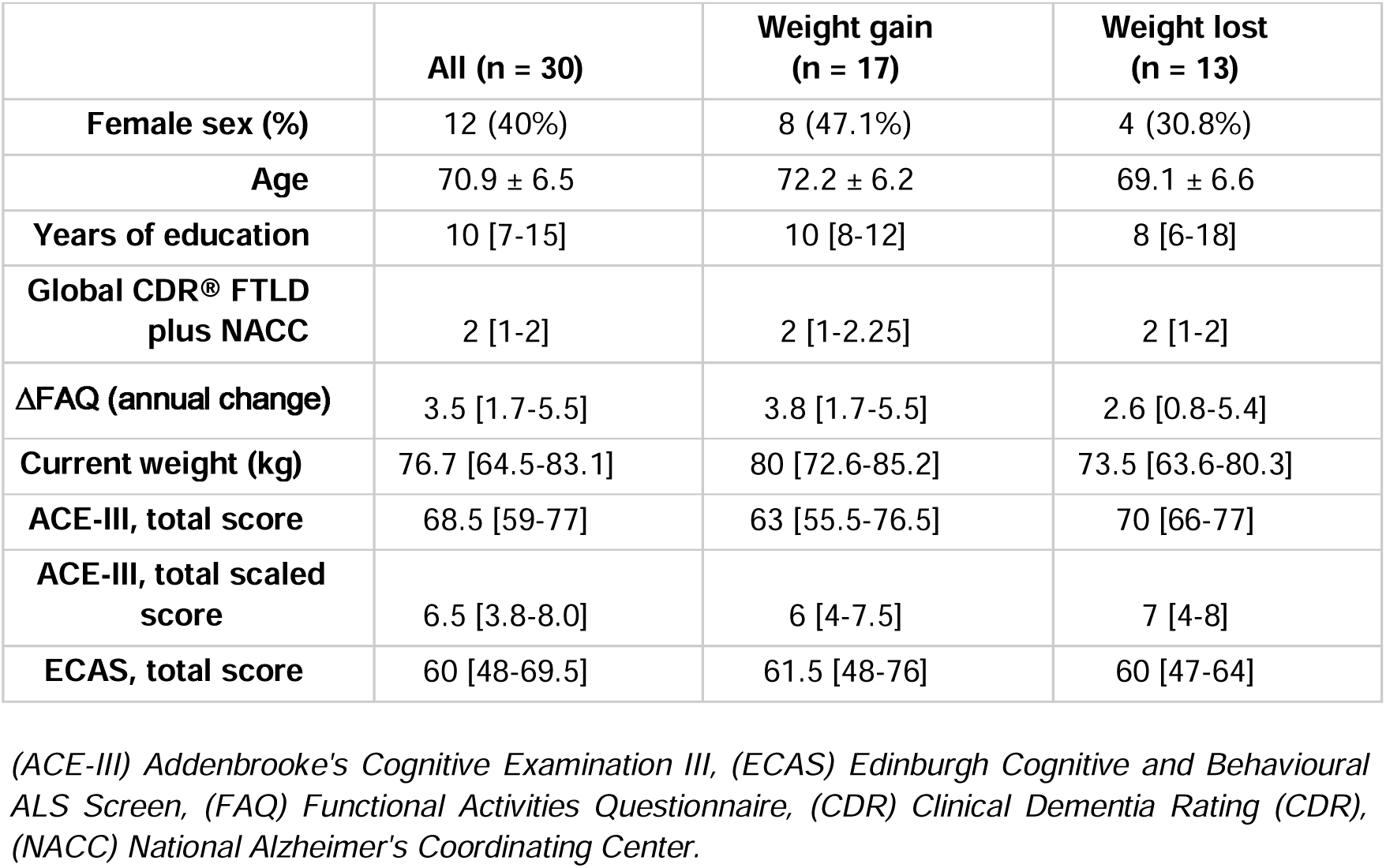
Demographic and neurological data for the patient cohort.

All FTD patients exhibited comparable baseline Clinical Dementia Rating (CDR) scores (Table 1), with some differences in their functional progression. Notably, the annual delta of the Functional Activities Questionnaire (FAQ)—which evaluates instrumental activities of daily living—tended to be lower in patients with weight loss compared to those who gained weight (Fig. 1c). General cognitive performance and fronto-executive function, assessed using the Addenbrooke’s Cognitive Examination (ACE) and the Edinburgh Cognitive and Behavioural ALS Screen (ECAS), were comparable between FTD patients who gained weight and those who lost weight (Table 1 and Fig. S1c, d). Similarly, serum levels of total cholesterol, LDL cholesterol, and triglycerides showed no association with weight changes (Fig. S1e-g). Together, these findings indicate that changes in body weight are a poor surrogate marker of metabolic dysfunction in FTD, as they do not reflect either cognitive status or systemic lipid alterations in this cohort.

We next investigated whether serum triglyceride, total cholesterol, and LDL levels were associated with neuropsychological measures of global cognition and fronto-executive function in bvFTD patients stratified by weight trajectory. In this cohort, serum triglycerides and LDL exhibited consistent inverse correlations with performance in tests assessing processing speed, working memory, verbal fluency, and executive function (Fig. 1d-f and Supplementary Table 1). Overall, higher circulating triglyceride and LDL levels were associated with poorer cognitive performance.

Overall, metabolic alterations and blood lipid alterations were observed in most patients regardless of them gaining or losing weight, and in particular blood lipids dysregulation were associated with worse cognitive performance.

Elucidating the molecular mechanisms that couple lipid metabolism to neuronal dysfunction will therefore be essential for understanding the biological basis of this metabolic vulnerability and for developing targeted nutritional and therapeutic strategies tailored to the metabolic profile of individual patients.

### 2. Life-long high-fat diet worsened metabolic and cognitive alterations in TDP-43 mutant mice

Prompted by these observations in bvFTD patients, we used a physiological TDP-43 knock-in mouse model to investigate the mechanistic relationship between metabolic vulnerability and disease severity. The TDP-43-M323K mouse model carries a point mutation that arose from a chemical mutagenesis experiment, this changes base pair c.968T>A of the mouse *Tardbp* gene resulting in the TDP-43 M323K missense mutation towards the C-terminus of the protein. This mouse strain exhibits neurodegeneration (18) and cognitive impairment (17) together with systemic metabolic disturbances driven, at least in part, by dysregulation of cholesterol metabolism (8). Based on our hypothesis, we anticipated that TDP-43-induced brain lipid dysregulation would elicit compensatory behavioural responses, leading to changes in food intake and dietary preferences. Thus, we first measure the food consumption and food preferences in mice with a physiological TDP-43 alteration compared to their littermate controls. *TDP-43^M323K/M323K^* mice had a non-significant trend towards increased eating, consistent with some patients (Fig. S2a, b). However, mutant mice had a food preference for a fattier and hypercaloric type of food (with 18% kcal from fat), compared to their littermates (standard chow with 12% kcal from fat) (Fig. 2a and Fig. S2c). Thus, this model provided a valuable platform to investigate the molecular consequences of dietary metabolic modulation in a brain already affected by disease with underlying potential compensatory mechanisms, as well as its longitudinal impact on cognitive function.

**Figure 2.**
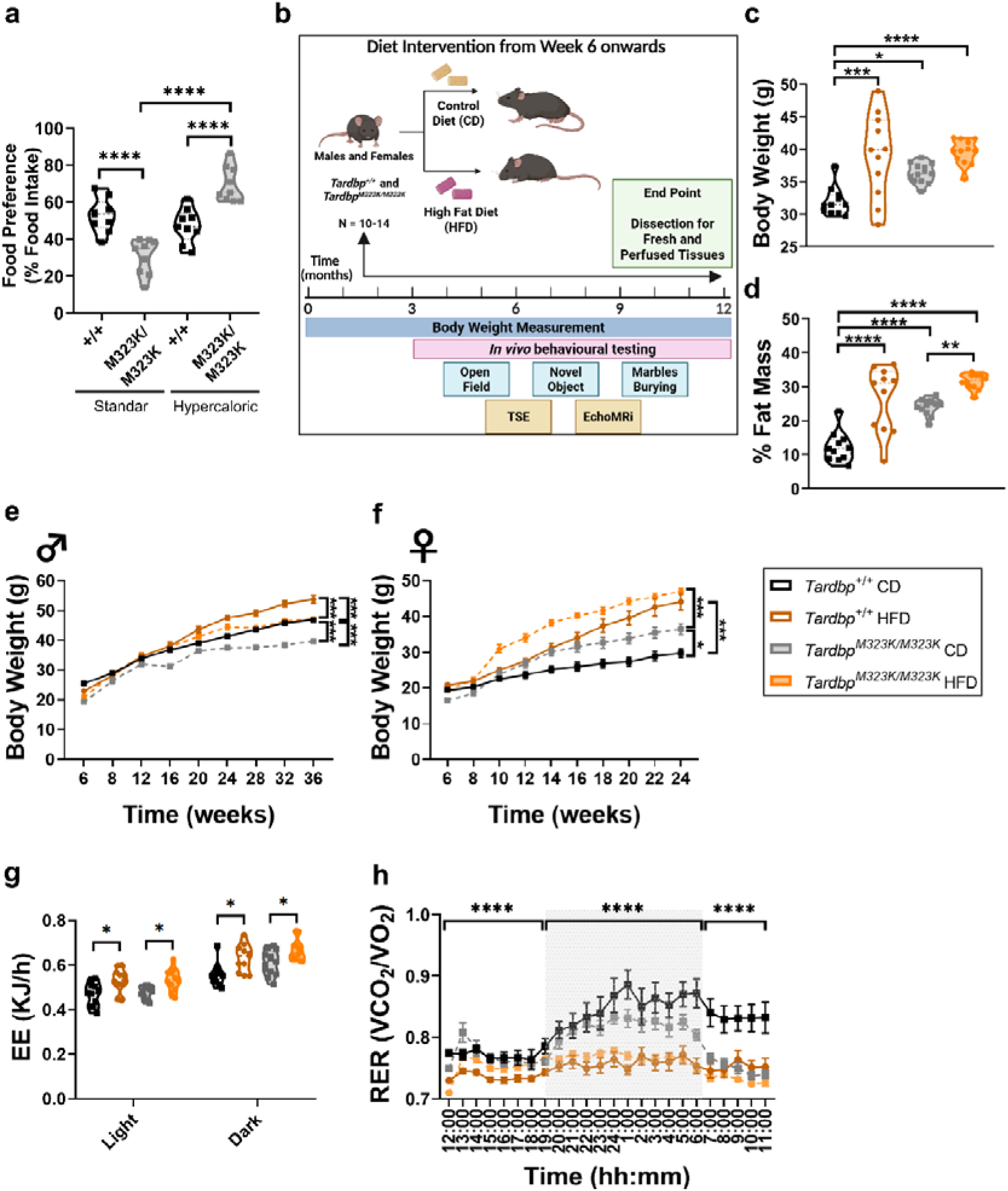
Metabolic and Cognitive impact of High Fat Diet in a TDP-43 proteinopathy model. **a.** Food preference in 9 month-old males, n: Tardbp^+/+^ = 10, Tardbp^M323K/M323K^ = 9. **b**. Experimental design of the dietary intervention**. c**. Body weight 3 months males; n: Tardbp^+/+^ CD = 11, Tardbp^M323K/M323K^ CD = 13, Tardbp^+/+^ HFD = 11, Tardbp^M323K/M323K^ HFD = 12 in males. **d**. Fat mass MRi males; n: Tardbp^+/+^ CD = 11, Tardbp^M323K/M323K^ CD = 13, Tardbp^+/+^ HFD = 11, Tardbp^M323K/M323K^ HFD = 12 **e**. Body weight curve in males; n: Tardbp^+/+^ CD = 10, Tardbp^M323K/M323K^ CD = 9, Tardbp^+/+^ HFD = 14, Tardbp ^M323K/M323K^ HFD = 14. **f**. Body weight in females; n: Tardbp^+/+^ CD = 12, Tardbp^M323K/M323K^ CD = 12, Tardbp^+/+^ HFD = 12, Tardbp^M323K/M323K^ HFD = 12 **g**. Indirect calorimetry, energy expenditure measurements; n: Tardbp^+/+^ CD = 11, Tardbp^M323K/M323K^ CD = 13, Tardbp^+/+^ HFD = 11, Tardbp^M323K/M323K^ HFD = 12. **h**. Indirect calorimetry, respiratory exchange ratio (RER); n: Tardbp^+/+^ CD = 11, Tardbp^M323K/M323K^ CD = 13, Tardbp^+/+^ HFD = 11, Tardbp^M323K/M323K^ HFD = 12.

Therefore, *TDP-43^+/+^* and *TDP-43^M323K/M323K^* mice were subjected to lifelong diets; either a control diet (**CD**) (10% of total energy from fat, 20% from protein, and 70% from carbohydrates) or a high-fat diet (**HFD**) (45% of the total energy from fat, 20% from protein, and 35 % from carbohydrates), starting at six weeks of age. Animals were followed longitudinally until 9 months (males) or 12 months (females), recording data for body weight, metabolic parameters, and behavioural performance, with tissues collected at the study endpoint for molecular analyses (Fig. 2b).

At an early time (3 months of age), only 6 weeks after the diet changed, the HFD induced an increase in body weight proportionally in both wild-type and in the mutant mice (Fig. 2c), mainly by a marked increase in fat mass (Fig. 2d), demonstrating the development of a robust obesogenic phenotype. Surprisingly, mutant male mice exhibit lower weight with age compared with control wild-type littermates, regardless of the diet, confirming the progressive metabolic dysregulation effect of the TDP-43 alteration which could not be compensated by the diet (Fig. 2e). This switching effect with age was not as clearly observed in mutant females (Fig. 2f). Interestingly, regardless of the mice gaining or losing weight, the TDP-43-M323K mutation induced higher systemic energy metabolism, measured by indirect calorimetry in a 24-hour metabolic cage, observed in the dark-active phase, and these differences were increased by the high-fat feeding, in both males and females (Fig. 2g and Fig. S2d). Similarly, the mutation induced increased lipid utilization as an energy substrate (measured by RER, in the dark phases), suggesting altered metabolic flexibility in mutant animals, which was further exacerbated in all mice by the HFD (Fig. 2h and Fig. S2e).

These data indicate that TDP-43 dysfunction induced a metabolic vulnerability that was exacerbated by a high fat diet. Importantly, this phenotype was largely independent of body weight trajectory, reinforcing that weight gain and weight loss was not representative of the underlying metabolic dysregulation.

### 3. High-fat diet exacerbates cognitive deficits in TDP-43 mutant mice

Thus, by modelling the effect of the metabolic dysregulation observed in some FTD patients, we next sought to determine whether this lifelong HFD could modify the behaviour of mice harbouring the TDP-43 M323K mutation. Longitudinal behavioural testing revealed progressive alterations in exploratory and cognitive performance. In the marble burying test, *Tardbp^M323K/M323K^*mice displayed a significant and progressive reduction in the percentage of buried marbles compared with wild-type controls. Importantly, HFD markedly exacerbated this behavioural alteration, both in males and in females (Fig. 3a and Fig. S3a). Short-term memory was assessed using the novel object recognition (NOR) task. Again, *Tardbp^M323K/M323K^* mice exhibited a progressive decline in discrimination ratio which was significantly aggravated in the mutants with HFD, suggesting that metabolic stress accelerates cognitive decline associated with TDP-43 dysfunction (Fig. 3b and Fig. S3b). In the open field test, *Tardbp^M323K/M323K^*mice showed reduced total distance travelled compared with wild-type animals, together with changes in center exploration. Again, these deficits were more pronounced in HFD-fed mutant mice (Fig. 3c and Fig. S3c).

**Figure 3.**
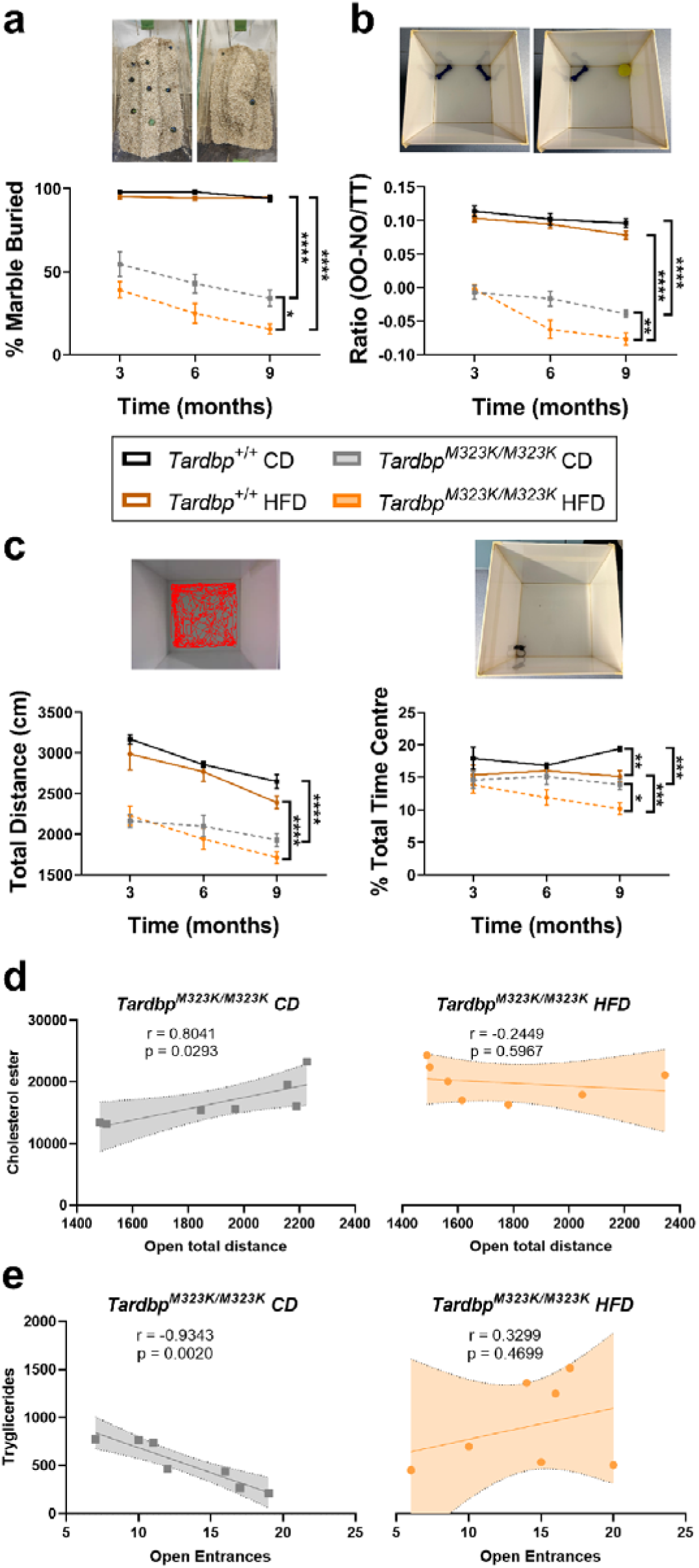
Lifelong high fat diet effects on the cognitive induced by the Tardbp ^M323K/M323K^ mutation in males. **a**. Marble burying test in males up to 9 months of age. **b**. Novel Object Recognition (NOR) test in males up to 9 months of age. **c**. Open field test in males up to 9 months of age, showing total distance travelled and time spent in the center. (a-c: n: Tardbp^+/+^ CD = 10, Tardbp^M323K/M323K^ CD = 9, Tardbp^+/+^ HFD = 14, Tardbp^M323K/M323K^ HFD = 14 in males.). **d.** Correlation between Cholesterol ester levels and Open total distance. **e.** Correlation between Triglycerides levels and Open Entrances. d-e: n: Tardbp^+/+^ CD = 7, Tardbp^M323K/M323K^ CD = 7, Tardbp^+/+^ HFD = 7, Tardbp^M323K/M323K^ HFD = 7 in males.

We looked at the levels of triglycerides and cholesterol ester in serum in relation to the behavioural data. Similar to findings from FTD patients, the different diets had an effect on the correlations between lipid levels and behavioural performance in mutant mice (Fig. 3d and Fig. S3d).

Thus, mirroring the clinical observation that systemic metabolic alteration was associated with worse disease progression in FTD patients, a high fat diet exacerbated behavioural impairments in TDP-43 mutant mice, supporting the notion of an underlying metabolic vulnerability.

### 4. Multiomics analyses revealed altered cholesterol and fatty acid metabolism pathways in TDP-43 mutant brain cortex

In order to identify the molecular mechanisms underlying the maladaptive metabolic pathways triggered by the interaction between TDP-43 dysfunction and a lipid-rich diet, we next examined the brains of these mice at 9 months of age and conducted an integrated multi-omics analysis of the brain frontal cortex.

First, we conducted a transcriptomic analysis, via bulk RNA sequencing of the frontal cortex from 9-month-old male mice and identified 223 differentially expressed genes (DEGs) *p* adj < 0.05 comparing *TDP-43^M323K/M323K^*against *Tardbp^+/+^* HFD-fed mice (Fig. S4a), which was comparable in magnitude to the 226 DEGs found when evaluating the effect of the mutation alone under a control diet (8). However, the HFD specifically altered the fatty acid biosynthetic process, suggesting a diet-induced metabolic shift (Fig. S4b). To assess baseline differences between wild-type (WT) animals under both diets, we identified 523 differentially expressed genes (DEGs); however, pathway enrichment analysis revealed no significantly enriched pathways. To further elucidate how HFD modifies the mutational phenotype, we compared mutant mice under both feeding regimes (*TDP-43^M323K/M323K^* CD vs. *TDP-43^M323K/M323K^* HFD), identifying 734 DEGs, *p* value < 0.05 (Fig. 4a). Pathway analysis showed that the HFD in mutant mice primarily deregulated cholesterol and isoprenoid biosynthesis, neuron branching maturation, and cell chemotaxis (Fig. 4b). Knowing the role of TDP-43 alteration in cholesterol biosynthesis and transport genes in the brain, we looked at how an external lipid-rich diet was affecting those already compromised lipid pathways in the brain. Heatmap analysis of cholesterol-related genes revealed a marked exacerbation of the transcriptional dysregulation of cholesterol biosynthetic pathways in HFD-fed mutant mice, while genes involved in cholesterol transport remained comparatively unaffected (Fig. 4c). Thus, we expected an increment in the CE levels in the brain in the *TDP-43^M323K/M323K^* mice with HFD. We quantified cholesterol ester levels in the frontal cortex of the mice. While cholesterol ester levels were significantly increased in mutants compared to controls across both dietary conditions, no significant differences were observed due to the diet (Fig. 4d). Accordingly, the amount of accumulated lipid droplets in the frontal cortex were proportional to the levels of CE detected, mostly increased in the mutants regardless of the diet (Fig. S4c).

**Figure 4.**
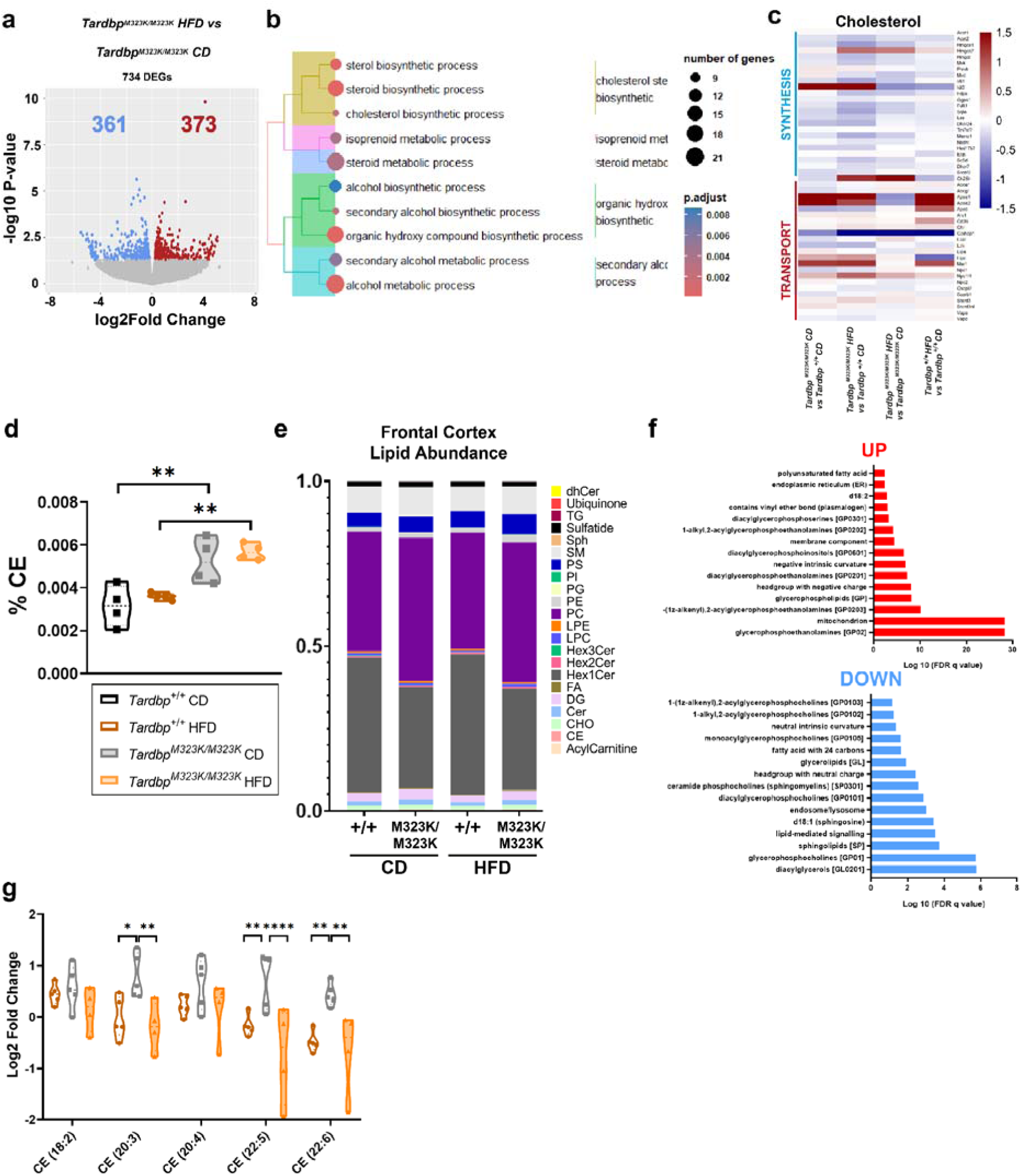
Global alterations induced by the Tardbp^M323K/M323K^ mutation and modulated by High Fat Diet (HFD) in the frontal cortex. **a**. Volcano plot of the 734 DEGs between the Tardbp^M323K/M323K^ HFD vs Tardbp^M323K/M323K^ CD comparison. **b.** Over-representation analysis (ORA) of deregulated genes. **c**. Heatmap of cholesterol–related genes in the frontal cortex of the mice. **d**. Cholesterol ester in the frontal cortex of the mice. **e**. A stacked bar chart showing the relative abundance of all detected lipid classes. **f**. Lipid Ontology (LION) enrichment analysis with upregulation and downregulation **g**. Fold change in CEs with indicated number of double bonds (unsaturation) in each group versus Tardbp^+/+^ CD. a-c: Experimental n = 4 for all groups. e-g: Experimental n = 5 for all groups except Tardbp^M323K/M323K^ HFD n = 4.

To complement these transcriptomic findings, we conducted a lipidomic profiling of the contralateral frontal cortex, where the TDP-43 mutation emerged as the primary driver of lipidomic levels variance (Fig. 4e). Notably, sphingomyelin (SM) levels showed a significant reduction in HFD-fed *TDP-43^M323K/M323K^* mice compared to HFD-fed *Tardbp^+/+^*, a difference absent under control diet conditions. Thus, the mutation, rather than the diet, accounted for most of the alterations at the level of the major lipid classes (Fig. 4e). Still, the lifelong high fat diet induced subtle changes in several lipids, sufficient to perform a pathway enrichment analysis comparing the mutants under HFD with the mutants on the control diet. The HFD promoted the upregulation of lipids involved in mitochondrial energy metabolism and reduced lipids involved in signalling (Fig. 4f). Since the level of unsaturation in the fatty acid chains has a major influence in those two processes (utilization in mitochondrial energy metabolism and signalling), we had a closer look at the level of unsaturation of cholesterol in the frontal cortex of these mice. In the control *Tardbp^+/+^* mice, the HFD had slight effect on reducing the level of unsaturated amino acids chains of CE and displayed a marked reduction in highly polyunsaturated cholesterol esters (20:3, 22:5, and 22:6; (Fig. 4g)). indicative of altered lipid esterification and broader dysregulation of lipid homeostasis. Such changes may promote a more rigid and pro-inflammatory membrane environment, potentially increasing neuronal vulnerability.

An integrative multi-omics analysis was performed using the DIABLO framework identified a highly coordinated transcriptome–lipidome network that discriminated mutant mice fed a high-fat diet from those maintained on the control diet (|r| ≥ 0.90). The analysis pointed towards a global repression of mitochondrial oxidative metabolism and sterol biosynthesis, evidenced by strong inverse correlations between key regulatory genes, including *Acadm* and *Cyp27a1*, and the accumulation of phosphatidylcholine (PC) species and free fatty acids (FFAs). Several lipid metabolism-related genes, including *Acadm*, *Cyp27a1*, *Adipor2*, *Capn2*, *Elovl1* and *Pld6*, exhibited strong coordinated associations with a common set of lipid species, defining a distinct molecular module. Most of these genes were negatively correlated with multiple phosphatidylcholine (PC) species and free fatty acids (FFAs), while showing positive correlations with specific sphingolipids, including Hex1Cer (d18:1/24:0), Hex1Cer (d18:1/22:0) and SM (41:2), likely related to cellular stress and inflammatory responses to the HFD. Conversely, *Inpp4a* displayed the opposite correlation pattern, suggesting the presence of a distinct transcriptome–lipidome module associated with the metabolic remodelling induced by the high-fat diet (Fig. S4d). Collectively, the transcriptomic, lipidomic, and integrative multi-omics analyses revealed that the interaction between TDP-43 dysfunction and a high-fat diet primarily disrupts three interconnected biological processes in the frontal cortex: sterol and cholesterol metabolism, mitochondrial oxidative and fatty acid metabolism, and lipid-mediated signalling pathways. These genotype-diet-specific alterations were accompanied by the accumulation of specific phosphatidylcholine and free fatty acid species, reduced lipid unsaturation, and coordinated transcriptional changes indicative of impaired metabolic adaptation. Given the well-established role of TDP-43 as a regulator of RNA processing, these findings raised the possibility that the profound metabolic remodelling induced by the HFD could also modify the splicing activity of TDP-43, thereby affecting downstream neuronal pathways beyond metabolism itself.

### 5. High-fat diet reprograms TDP-43 splicing activity toward synaptic dysfunction in the mutant frontal cortex

To further look at why the adaptive metabolic shift triggered by the HFD was impaired in the mutant TDP-43 brains, we specifically looked at the particular metabolic processes regulated by TDP-43 through its role in the regulation of RNA metabolism. Thus, we used the RNA sequencing data from the frontal cortex of the mice in all the experimental groups.

First, we looked at RMAPS analysis of *Tardbp*, which revealed a diet-dependent redistribution of *Tardbp* binding motifs, indicating that the metabolic state significantly modulates the splicing regulatory landscape of TDP-43. Under diet conditions, the mutation was associated with strong motif enrichment around alternatively spliced exons, characterized by predominant red peaks upstream and downstream of exons (Fig. 5a) suggesting that mutant TDP-43 preferentially promotes exon inclusion. In contrast, HFD exposure shifted this pattern toward a predominance of blue peaks (Fig. 5b) consistent with a functional switch towards exon repression and exclusion. The altered distribution of TDP-43 motifs near splice sites may reflect changes in protein occupancy that could interfere with spliceosome assembly and efficient intron excision, potentially contributing to the exacerbated splicing defects observed under HFD conditions.

**Figure 5.**
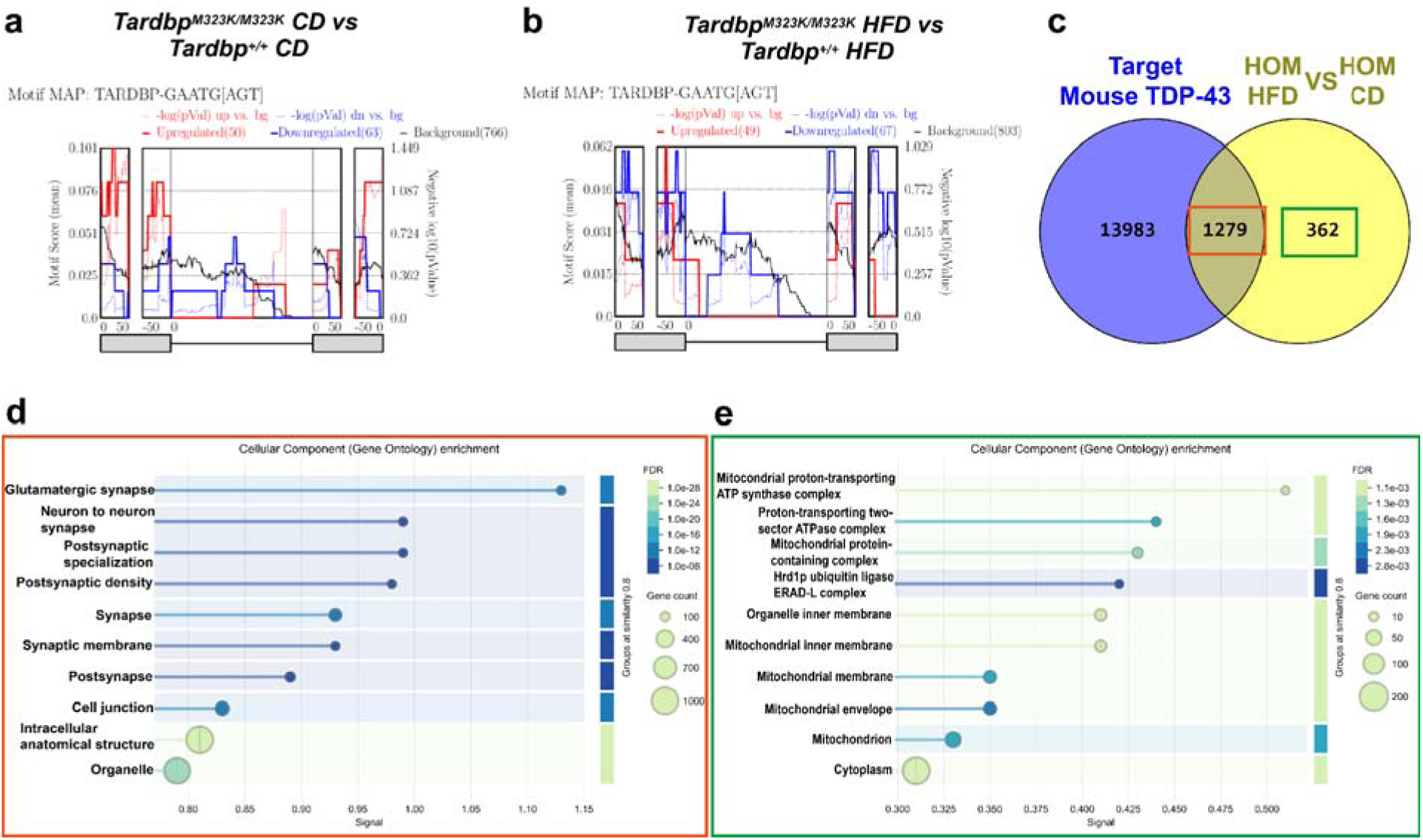
**a**. Distribution of TARDBP (GAATG[AGT]) binding motif enrichment around retained introns in mutant vs control under control **b**. Distribution of TARDBP (GAATG[AGT]) binding motif enrichment around retained introns in mutant vs control under high-fat diet. **c**. Venn diagram of the Tardbp^M323K/M323K^ CR vs Tardbp^M323K/M323K^ CD comparison showing TDP-43 target overlap. **d**. Gene Ontology enrichment of the 1279 TDP-43 target genes exhibiting at least one alternative splicing event in the cortex. **e**. Gene Ontology enrichment of the 1279 non-TDP-43 target genes exhibiting at least one alternative splicing event in the cortex. Experimental n = 4 for all groups.

To study the different effect of the HFD on the functionality of the wild-type and mutant TDP-43, we looked at the global spliceosome. The HFD impacted on the functional activity of the mutant TDP-43, observing a difference of 1660 events compared to the mutant mice fed with the control diet. From those, 1279 genes were identified as TDP-43 targets (Fig. 5c). The pathway analyses of those 1279 genes showed that the main pathways involved were related to postsynapse and cell junction in the frontal cortex (Fig. 5d). In contrast, the genes exhibiting changes in splicing events that were not targets of TDP-43 induced by the HFD were involved in the regulation of mitochondrial protein complex–related pathways (Fig. 5e), supporting the changes observed with the lipidomic data (Fig. 4d).

The particular interaction between the mutant TDP-43 specific targets and the modification induced by the HFD pointed, at least partially, towards modification at the synapses. A further modification at the synapses in the mutants with HFD could explain the enhanced cognitive impairment in these mice.

### 6. High-fat diet exacerbates brain dysfunction in TDP-43 mutant mice

Together, these multi-omics analyses revealed that HFD reprograms metabolic pathways and the RNA-processing functions of TDP-43 in the mutant brain, particularly affecting mitochondrial metabolism, sterol homeostasis and synaptic gene regulation. We next asked whether this molecular remodelling was reflected in brain function. To address this, we evaluated neuroinflammation, cerebral glucose metabolism by FDG-PET, and cortical neurotransmitter homeostasis.

Immunofluorescence analysis of the microglial marker IBA1 revealed a further significant increase in microglial activation in mutant mice compared with wild-type controls under the HFD (Fig. 6a). Similarly, staining for the astrocytic marker GFAP demonstrated increased astrocyte activation in response to HFD, which was particularly prominent in mutant animals (Fig. 6b). These results indicated that dietary lipid overload enhances neuroinflammatory responses in the context of TDP-43 pathology.

**Figure 6.**
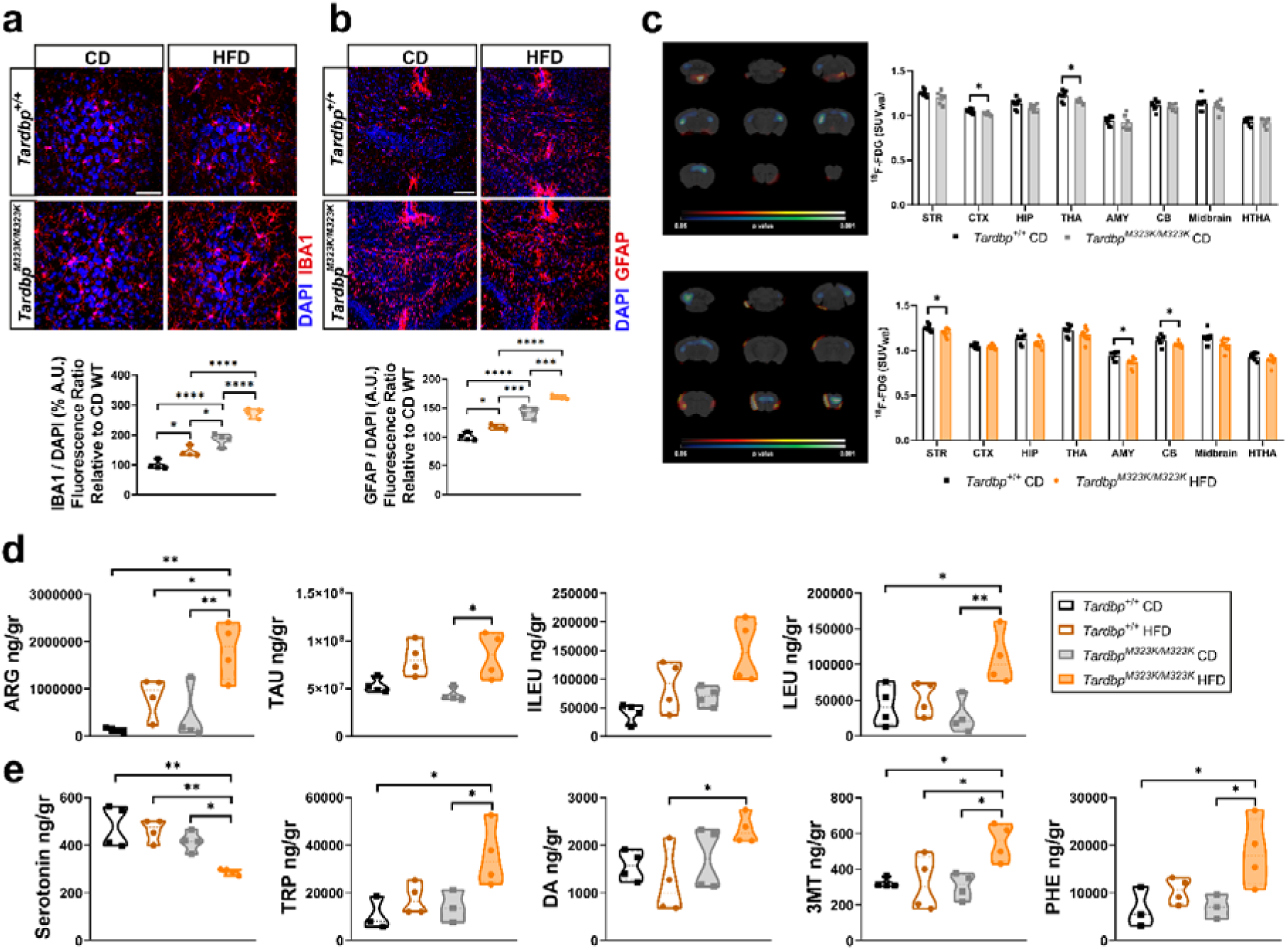
A high-fat diet induces functional changes in the mutant mice. **a.** Representative image and quantification of IBA1 immunostaining (IBA1 in red and Hoetsch in blue) in the frontal cortex of 9-month-old males. Scale bar= µm. **b**. Representative image and quantification of GFAP immunostaining (GFAP in red and Hoetsch in blue) in the frontal cortex of 9-month-old males. Scale bar= µm. **c**. Group average PET-glucose image and quantification. n: *Tardbp^+/+^* CD = 8, *Tardbp^M323K/M323K^* CD = 7, *Tardbp^M323K/M323K^* HFD = 8 in females. **d**. Neurotransmitter levels in the frontal cortex, including arginine (ARG), serotonin, taurine (TAU), isoleucine (ILEU), leucine (LEU). **e**. Neurotransmitter levels in the frontal cortex, including serotonin, tryptophan (TRP), dopamine (DA), 3-methoxytyramine (3-MT), and phenylalanine (PHE), expressed as ng/g. All samples correspond to males, and the experimental n for all groups = 4.s

To assess the brain functional consequences of those molecular changes, we performed Positron Emission Tomography with 18-Fluorodeoxyglucose (PET-^18-^FDG) in a selected group of female mice at 12 months of age (Fig. 6c). The PET signal was significantly reduced in the *Tardbp^M323K/M323K^* mutant mice relative to their wild-type controls in thalamus and cortex under control diet conditions, indicating localized hypometabolism. In contrast, in mutant mice fed with a HFD, these alterations were no longer observed in the thalamus and cortex, and significant reductions in PET signal emerged in the amygdala, cerebellum, and striatum, indicating a shift in the regional pattern of hypometabolism. Thus, despite some regional variability in non-significant trends, all statistically significant changes pointed toward a reduction in metabolic activity (Fig. 6c). These findings indicated a model in which the *Tardbp* mutation drives region-specific cerebral hypometabolism, while the regional effects were further modulated under metabolic stress conditions induced by a high-fat diet.

Finally, we further characterized the impact of the mutation and the HFD on cortical function by profiling cortical neurotransmitter levels (Fig. 6d, e). Targeted metabolomic profiling of the frontal cortex in 9-month-old mice revealed that HFD exposure in *Tardbp^M323K/M323K^* mice further disrupted metabolites linked to synaptic bioenergetic dysfunction. L-arginine (ARG) levels were markedly increased in HFD mice, suggesting activation of iNOS- and arginase-dependent microglial pathways and a state of cortical neuroinflammation driven by the combined effects of lipotoxicity and the TDP-43 mutation (19). Similarly, taurine (TAU) levels were selectively elevated in this group, more likely reflecting an adaptive response to oxidative stress. Furthermore, the branched-chain amino acids leucine (LEU) and isoleucine (ILEU) reached their highest concentrations in HFD-fed mutant mice. Those alterations could indicate impaired mitochondrial amino acid catabolism associated with brain insulin resistance and neuronal energy dysfunction (20).

Analysis of monoaminergic pathways revealed a marked disruption of the serotonergic system, characterized by increased tryptophan (TRP) levels together with a selective reduction in serotonin (5-HT) in HFD-fed mutant mice, indicative of an enzymatic bottleneck in Tryptophan Hydroxylase (TPH) activity (21), likely precipitated by the ATP deficit. In contrast, the dopaminergic pathway exhibited a signature of metabolic hyperactivity, with coordinated increases in phenylalanine (PHE), dopamine (DA), and 3-methoxytyramine (3MT), consistent with enhanced catecholamine synthesis and increased cortical dopamine turnover associated with chronic high-fat diet consumption (22).

Collectively, these findings indicated that TDP-43 dysfunction was associated with persistent cerebral hypometabolism, which was further exacerbated by a high-fat diet. This metabolic impairment was accompanied by cortical metabolic remodelling, including altered amino acid and monoamine metabolism, together with increased neuroinflammatory responses, consistent with impaired synaptic bioenergetics and neurotransmission.

## Discussion

Altered eating behaviour is among the earliest and most characteristic clinical manifestations of behavioural variant frontotemporal dementia (bvFTD) (23,24). Over the past two decades, these behavioural changes have been primarily interpreted as consequences of frontal and hypothalamic degeneration, supported by neuroimaging and neuropathological studies demonstrating dysfunction of neural circuits regulating reward processing, satiety and food motivation (25–27). However, an increasing number of clinical and experimental studies have also identified profound systemic metabolic alterations in FTD, and particularly bvFTD, including increased caloric intake, selective preferences for sweet and high-fat foods, weight gain, increased adiposity, visceral fat accumulation, dyslipidaemia and insulin resistance (28,29). Despite these observations, the relationship between metabolic dysfunction and the behavioural phenotype of bvFTD remains poorly understood. This central question remains open, and the direct empirical evidence is limited, whether altered eating behaviour simply reflects neuronal dysfunction or could also represent an adaptive response to an underlying metabolic disturbance.

Our findings support the latter possibility and provide evidence that the biological consequences of increased lipid availability depend less on nutrient exposure itself than on the metabolic resilience of the diseased brain. By combining clinical observations with mechanistic studies in a physiological *Tardbp* knock-in model, we propose that altered eating behaviour may initially represent a compensatory response to impaired cerebral metabolic homeostasis. However, in the context of TDP-43 dysfunction, this adaptive response ultimately fails because the metabolically vulnerable brain cannot appropriately remodel its metabolic and RNA regulatory programmes in response to chronic lipid stress.

Although weight gain has often been considered a characteristic metabolic feature of bvFTD, clinical studies have consistently reported considerable heterogeneity between patients (13,30).

Clinical imaging work further shows that patients with bvFTD can have increased android fat deposition and visceral adipose tissue despite BMI matching, indicating that body weight alone is an incomplete marker of metabolic status (31). Recent metabolomic and lipidomic studies further support this view by demonstrating widespread alterations in systemic lipid metabolism that correlate more closely with disease severity than with anthropometric measures (32–34). Several of these circulating lipid signatures correlate more closely with disease severity, prognosis, cognition, and functional decline than with body size alone (35) Our clinical observations extend these findings by showing that circulating lipid abnormalities, rather than body weight itself, were associated with poorer cognitive performance. Together, these data reinforce the concept that metabolic dysfunction represents a fundamental component of bvFTD biology rather than merely a secondary consequence of altered eating behaviour.

Experimental models have established that TDP-43 dysfunction disrupts multiple aspects of cellular metabolism, including mitochondrial activity, cholesterol homeostasis and glucose utilization (10,36–38). TDP-43 also regulates brain cholesterol homeostasis more directly through SREBP2/SREBF2-dependent pathways in oligodendrocytes and other experimental systems, with downstream effects on myelination and brain cholesterol content (8,37,39). However, most previous studies have examined these alterations under basal conditions, providing limited insight into how metabolically vulnerable brains respond to environmental metabolic challenges. Using chronic high-fat feeding as a metabolic stressor, trying to partially mimic the effect of altered eating behaviour in FTD patients, our study demonstrates that increased lipid availability does not restore metabolic homeostasis but instead unmasks a profound loss of metabolic flexibility. This observation shifts the interpretation of high-fat feeding from a simple dietary manipulation to a functional probe of cerebral metabolic resilience.

Mechanistically, the integrated multi-omics analyses revealed a coordinated metabolic rewiring upon the challenge of a high fat diet. One of the principal advances of the present work is the integration of transcriptomic, lipidomic and RNA splicing analyses within the same experimental framework. Although disrupted cholesterol metabolism, mitochondrial dysfunction and defective RNA processing have each been independently implicated in TDP-43 proteinopathies, these mechanisms have generally been investigated separately. TDP-43 loss of function is now strongly linked to RNA-processing defects, including cryptic exon inclusion, altered exon usage, and cryptic alternative polyadenylation across iPSC-derived neurons, postmortem ALS/FTD tissue, and presymptomatic disease states (40–42). In parallel, TDP-43 dysfunction has been linked to cholesterol dysregulation and broader lipid pathway abnormalities, including reduced endogenous cholesterol synthesis, altered transport programs, cholesterol-related lipid species changes, and lipid droplet accumulation in mouse models and FTLD-TDP tissue (8,43). Mitochondrial and oxidative bioenergetic defects are also well supported, with integrated transcriptomic-proteomic studies identifying early compensatory signatures in fatty-acid beta-oxidation and mitochondrial protein translation, and mechanistic work showing that TDP-43 aggregation perturbs nuclear-encoded mitochondrial programs and amplifies oxidative stress (44–46). Our data instead suggest that chronic metabolic stress coordinates these apparently distinct pathological processes into a unified molecular response. The convergence of transcriptional, lipidomic and splicing alterations on pathways regulating sterol metabolism, oxidative bioenergetics and synaptic organization indicates that metabolic stress reorganizes interconnected cellular networks rather than simply exacerbating isolated defects.

Aberrant RNA processing is widely recognized as a defining feature of TDP-43 proteinopathies, because loss of nuclear TDP-43 disrupts hundreds of splicing events and alters RNA programs enriched for synaptic and neuronal transcripts (47,48). TDP-43-dependent mis-splicing is now established in disease-relevant targets such as UNC13A, STMN2, and KCNQ2, and broader studies show that these changes extend to multiple genes controlling synaptic transmission and membrane excitability (49–51). Whether the RNA-regulatory activity of TDP-43 is itself influenced by metabolic state has remained comparatively unexplored, even though TDP-43 dysfunction is also linked to bioenergetic failure, lipid metabolic rewiring, and disturbed cholesterol homeostasis. This gap matters because cell context and stress state can change TDP-43-dependent RNA processing, and stress-induced nuclear TDP-43 condensation has already been shown to cause transient loss of splicing function(52,53).

The molecular consequences unmasked by the HFD identified by our multi-omics analyses were paralleled by cerebral hypometabolism, neurotransmitter imbalance and enhanced glial activation. These observations are consistent with a growing body of literature indicating that impaired bioenergetics constitutes an early and sustained feature of FTD (54–58). Importantly, our data integrate these previously independent observations into a common framework in which defective metabolic adaptation links altered lipid utilization, synaptic dysfunction and neuroinflammatory responses (59). Rather than representing parallel pathological processes, these alterations appear to reflect different manifestations of a common failure to preserve neuronal metabolic homeostasis under chronic metabolic stress.

Collectively, our findings support a reinterpretation of the metabolic phenotype of bvFTD. Rather than viewing altered eating behaviour solely as a behavioural consequence of neurodegeneration, our data suggest that it may also reflect an attempt to compensate for an underlying metabolic vulnerability. Although this adaptation may initially be beneficial, the presence of TDP-43 dysfunction appears to compromise the capacity of the brain to respond appropriately to sustained lipid excess, ultimately promoting coordinated alterations in metabolism, RNA processing and neuronal function. More broadly, these findings highlight metabolic resilience as a previously underappreciated determinant of disease progression and suggest that future metabolic interventions should aim not simply to increase energy availability but to restore the capacity of vulnerable neuronal networks to adapt to metabolic stress.

### Limitations of the study

This study has several limitations that should be considered when interpreting the findings. First, the relatively small cohort of patients with frontotemporal dementia (FTD) limits the statistical power to detect subtle associations and may reduce the generalizability of the observed correlations. Validation in larger, independent cohorts will be necessary to confirm these findings and further explore the relationships identified.

Second, neuropathological confirmation was not available for the patients included in this study. Consequently, it was not possible to determine the underlying molecular pathology, such as TDP-43 or tau pathology, within the FTD cohort. Given the biological heterogeneity of FTD, this limitation may have introduced variability into the analyses. Future studies incorporating neuropathological diagnosis or validated in vivo biomarkers of protein pathology will be essential to establish whether the observed findings are specifically associated with TDP-43 proteinopathy or are shared across different FTD subtypes.

### Conclusions

Our study suggests that the biological consequences of altered lipid availability depend on the metabolic state of the brain. By integrating clinical observations with mechanistic studies in a physiological TDP-43 knock-in model, we show that pre-existing metabolic vulnerability profoundly alters the cerebral response to chronic metabolic stress, resulting in coordinated metabolic, molecular and functional abnormalities. These findings provide a conceptual framework in which metabolic resilience emerges as a key determinant of disease progression and highlight the need to consider the underlying molecular pathology when designing metabolic and nutritional interventions for neurodegenerative diseases.

## Methods

### 1. Patients

Clinical and neuropsychological data were obtained from a cohort of patients with behavioural variant of frontotemporal dementia. This is a secondary analysis of data from a previously established cohort of patients with behavioural variant frontotemporal dementia (bvFTD) recruited at the Cognitive Neurology Unit of Hospital Clínico San Carlos (Madrid, Spain). Cases were included in the present analysis if they fulfilled the following criteria: (1) consensus criteria for probable bvFTD (60); (2) negative biomarkers for Alzheimer’s disease, determined by cerebrospinal fluid analysis, amyloid PET, or plasma p-tau217; and (3) absence of severe cerebrovascular disease (Fazekas grade 3) or any other significant structural brain lesion on neuroimaging.

All participants, or their legally authorized representatives when appropriate, provided written informed consent in accordance with institutional ethical standards and applicable regulations. Clinical trial number: not applicable.

Sociodemographic data, including sex, age, and years of formal education, were recorded. Body weight was measured at the time of assessment, and participants were asked to report their most stable body weight during the five years preceding symptom onset. When patients were unable to provide this information reliably, it was obtained from a caregiver or close relative. Based on the difference between current and premorbid body weight, patients were classified into two groups whether they gained or lost weight. Cognitive assessment included the Spanish versions of two screening instruments for bvFTD: the Addenbrooke’s Cognitive Examination III (ACE-III) (61) and the Edinburgh Cognitive and Behavioural ALS Screen (ECAS) (62). Although the ECAS was originally developed for patients with amyotrophic lateral sclerosis (ALS), it was included because it provides a more comprehensive assessment of executive functions than the ACE-III. In particular, whereas the ACE-III Attention domain combines orientation, immediate verbal memory, and a single serial subtraction task, the ECAS incorporates multiple executive function tasks together with a brief assessment of social cognition, a domain that is characteristically impaired in bvFTD, even during the prodromal stages of the disease (23). The ECAS has been shown to be a valid cognitive screening instrument for bvFTD (63). Because the two instruments share several subtests, they were administered during the same visit, and overlapping items were completed only once to minimize practice effects. Version A of the ECAS was used in all participants.

The ACE-III raw scores were converted into age- and education-adjusted scaled scores, according to the published normative data for the Spanish population (64). These scaled scores range from 2 to 18, with a score of 10 corresponding to the 50th percentile. Conversely, normative data for the Spanish version of the ECAS are not currently available. Therefore, ECAS results are presented as unadjusted scores, and age and years of education were included as covariates in the statistical analyses to account for their effects.

Finally, fasting venous blood samples were collected for the analysis of metabolic parameters, including total cholesterol, HDL cholesterol, LDL cholesterol (calculated using the Friedewald equation), triglycerides, prealbumin, glycated haemoglobin (HbA1c%), and the homeostatic model assessment for insulin resistance (HOMA-IR).

### 2. Animals

We use the *Tardbp^M323K/M323K^* line, maintained in a constant F1 DBA/2J-C57BL6/J hybrid background, as previously described (Godoy et.al 2024). For all the experiments we used the wild-type (*Tardbp ^+/+^*) and the homozygous littermates (*Tardbp ^M323K/M323K^*), and both sexes. Mice were kept in auto-ventilated cages with free access to water. Mice were kept under constant conditions with a regular 12 h light and dark cycle, temperature of 21 ± 2 ◦ C and humidity 55 ± 10%. Mice were housed in same sex cages of up to five mice. Cages contained aspen chips bedding (Datesand), shredded paper and a rodent tunnel for enrichment. The animals were always kept in their cages except in the case of a behavioural test. Isolation time was kept as short as possible so as not to affect the results of other behavioural tests. All mice were maintained and studied according to international, national and regional guidelines. The study was approved by the local ethical committees of animal care and use of the Hospital Clínico San Carlos (Proex 222.7/24 and 258.4/20) in accordance with the European and Spanish regulations (2015/39/EU and RD 53/2013).

At weaning, all mice were genotyped by PCR from an ear biopsy, using a multiplexed PCR assay. Forward primer: 5’-TCTGCAACGCTGCCTGAG-3’ and reverse primer: 5’-AGCTGGCTTGGGAAATAACCA-3’. Wildtype probe: 5’-CCGCAGCCATCATCG-3’ (5’ FAM-’3 BHQ1) and mutant probe: 5’-CGCAGCCTTCATC-3’ (5’ JOE - 3’ BHQ1). At 6 weeks of age months of age, all *Tardbp ^+/+^* and *Tardbp ^M323K/M323K^* mice were randomly assigned to one of the two feeding regimes: i/ Control diet (CD) (D12450H, Research Diets Inc., Brogaarden ApS, Danmark), or ii/ 45% High-Fat-Diet (HFD) (D12451, Research Diets Inc., Brogaarden ApS, Danmark) that link with control diet.

### 3. Phenotyping tests

All tests were conducted in both sexes separated, unless otherwise stated, and blind to the operator.

#### 3.1. Weights

Non-anaesthetised mice were weighed inside a beaker monthly and before the appropriate behavioural test. To correct for animal movements, an average weight was taken after five seconds.

#### 3.2. Marble burying test

Nine glass marbles were evenly spaced in rows of three in an IVC containing approximately 4 cm depth of aspen bedding. The mouse was placed in the cage, and IVC lids were positioned on top for 30 min. For the total score of buried marbles, the following scores were given <25% buried 0, around 50% buried 0.5 and >75% buried 1. The score was then normalized to the total number of marbles (9). Finally, the score was normalized to the average number of marbles buried by the wild-type at 3 months of age.

#### 3.3. Novel object recognition test

Initially, the mouse was habituated within the arena (methacrylate box of 40 cm × 40 cm). The habituation process lasted two days and the mouse spent 10 min in the box each day. On the third day, the test was carried out by measuring the time spent with each object. To do this, the mouse was placed in the box for 10 min with the two initial objects (two equal weights). One of the objects was removed (always the same one) and a novel object (a plastic lemon) was placed in the arena for the 5 min of the test. After 15 min, the mouse was removed and placed back into its home cage. The Novel Object box was cleaned before each run with 70% alcohol. To quantify the results, the following ratio was used: (time spent with the new object − time spent with the known object)/total time (300 s). Finally, the score was normalized to the ratio of time by the wild-type at 3 months of age.

#### 3.4. Open Field Test

Mice were placed in a 40 cm × 40 cm methacrylate box for behavioural testing. A camera was positioned above the arena to record the entire 10-minute session. Mice were allowed to explore freely for 10 minutes before being returned to their home cages. After completing the sessions and recording the videos, several measurements were performed. Total distance travelled was calculated automatically using EthoWatcher software. The videos were then manually reviewed twice: first to time the duration spent in the center of the arena, and second to count the number of entries into the center. The center was defined as a 30 cm × 30 cm square.

### 4. Positron Emission Tomography (PET)

Functional *in vivo* imaging scans were performed in groups of females at 12 months of age, divided in four groups with an experimental n between 6-9 per group. Imaging scans were performed on a hybrid positron emission tomography and computed tomography (PET/CT) scanner for small animals (Albira ARS, Bruker NMI, Valencia, Spain). Mice were intraperitoneally injected with the tracer ^18^F-Fluorodeoxyglucose (^18^F-FDG) (5.92 MBq / 200 μl saline solution, Curium Pharma, Madrid, Spain) for glucose metabolism evaluation. After 45 minutes of awake uptake, animals were anesthetized using isoflurane (2% in 100% oxygen) and placed in the PET/CT scanner with the brain in the central field of view. A 20-minute PET scan was performed followed by 10-minute CT acquisition. After image acquisition, PET scans were reconstructed using a maximum likelihood expectation maximization algorithm (MLEM, 12 iterations) and applying random, scatter, and decay corrections. CT images were reconstructed using filtered back projection. For image analysis, reconstructed CT images were manually registered to an in-house built magnetic resonance image (MRI) template (WT and TDP) (16165303) and subsequently spatial transformation was applied to the PET image corrected for whole brain uptake (SUV_WB_). A volume of interest (VOI)-template was then applied to the registered PET image and regional uptake was calculated. Additionally, registered PET images were subsequently used for parametric statistical analysis (SPM12; UCL, London, UK). Differences in ^18^F-FDG SU_WB_ between experimental groups were analyzed using a two-sample unpaired t-test using SPM12 under MATLAB software (The MathWorks Inc.). T-maps were loaded into PMOD, and a threshold was set for T-values corresponding to P = 0.05 (lower threshold) and P = 0.001 (upper threshold).

### 5. Neurotransmitters analyses

The dissected frontal cortex from one hemisphere from male mice of the 4 experimental groups (*Tardbp*^+*/*+^ and *Tardbp^M323K/M323K^* under two feeding regimes: CD-and HFD-fed), at 9 months of age (n = 4 per group, respectively) were sent to the Chromatography Service of Hospital Ramón y Cajal (IRICYS). To dissect the frontal cortex from the brain, the olfactory bulbs were first removed. A coronal cut was made at bregma 0, selecting all the anterior cortex carefully excluding the striatum. The two halves of the frontal cortex from the two hemispheres were then separated and one of them was used for the experiment. Cortical samples were sonicated in six volumes (weight/volume) of 0.4 N perchloric acid (PCA) for deproteinization and subsequently centrifuged at 10,000 × g at 4 °C for 20 min. Amino acids were determined by high-performance liquid chromatography (HPLC), as previously described (65). Fluorescence detection was performed using a Jasco FP-2020 detector at excitation and emission wavelengths of 240 nm and 450 nm, respectively. Amino acids were identified based on their retention times, and their concentrations were calculated by comparison with externally calibrated standard amino acid solutions (1.5 mM). Dopamine (DA) and its metabolites were measured from supernatants using high-performance liquid chromatography (HPLC) with an ESA Coulochem electrochemical detector, according to Menás protocol (66). Chromatographic conditions were as follows: an ACE 5 C18 column (150 mm × 4.6 mm; UK); a 0.1 M citrate/acetate buffer, pH 3.9, containing 10% methanol, 1 mM EDTA, and 1.2 mM heptane sulfonic acid. Detector voltages were set as follows: D1 = 10.05 V, D2 = 0.39 V, and guard cell = 10.40 V. Monoamine levels were identified based on retention times and quantified by comparison with externally calibrated standard solutions (0.6 mM).

### 6. Protein Study

#### 6.1. Immunostainings

Mice were terminally anaesthetised with 0.4 mg/kg of fentanest and 40 mg/kg of thiopental and cardiac perfused with the fixative 4% PFA without methanol in PBS 1×, pH 7.4. The brain was collected intact from the skull. The brain was postfixed in the same fixative overnight at 4 ◦C and kept on after PBS washes were cryopreserved by immersion in 30% sucrose for three consecutive days, changing every day. Tissues were dried and embedded in OCT using isopentane and dry ice, and frozen at − 20 ◦C until cut with a cryostat (Leica). Transverse sections at 35 μm were collected free-floating through the lumbar spinal cord and brain frontal cortex. Free-floating sections were then blocked with PTB (PBS 1×, Triton 0.02% and Bovine Serum Albumin (BSA)) and then incubated with the primary antibodies in PBS with Triton 0.02% (for the list of antibodies used, see Supplementary Table S2) and the secondary antibodies (Alexa-Fluor, ThermoFisher), and then mounted together with Hoechst stain to detect nuclei. Images were taken using a confocal microscopy Olympus Fluoview FV1000 with a version system 3.1.1.9.

#### 7. Imaging analysis

Image analysis and quantification were carried out using Fiji/ImageJ. Prior to measurement, the images were uniformly adjusted to ensure consistent thresholding across the samples. They were then converted into binary masks to isolate the signal of interest. The resulting segmented images were subjected to particle detection using the ‘Analyse Particles’ function with fixed size and circularity parameters applied to exclude background noise or artefacts. For each identified particle, the integrated density (IntDen), which reflects the sum of pixel intensities within the particle area, was extracted. The total signal intensity for each image was then derived by combining the particle count with the corresponding IntDen values. This provided a quantitative estimate of marker expression. This analytical approach was consistently applied to both IBA1 and GFAP immunoreactivity.

To quantify the number of lipid droplets (PLIN2) by cells, defined regions of interest (ROIs) or full-frame images were selected to ensure consistent sampling across sections. Following signal segmentation, cell-associated puncta were identified using the ‘Analyse particles’ function in Fiji/ImageJ. Detection parameters were adjusted to discriminate discrete objects approaching single-pixel resolution, while minimising background inclusion.

### 8. Targeted Lipidomics Analysis

Lipid extraction was conducted following the procedure previously described in (67). In brief, 10 µL of sample were mixed with 100 µL of a butanol:methanol solution (1:1, v/v) supplemented with 10 mM ammonium formate and the corresponding internal standards (Supplementary Table 3). Samples were vigorously vortexed and subsequently sonicated for 1 h at room temperature. After sonication, extracts were centrifuged at 14,000 × g for 10 min at 20°C, and the resulting supernatants were transferred into vials containing glass inserts for subsequent LC-MS analysis.

Lipidomic analyses were performed using an Ultivo Triple Quadrupole mass spectrometer coupled to an Agilent 1290 UHPLC system (Agilent Technologies, Barcelona, Spain). Chromatographic separation was achieved using a ZORBAX Eclipse Plus C18 column (2.1 × 100 mm, 1.8 µm particle size; Agilent Technologies) maintained at 60°C. Data acquisition was carried out in both positive and negative ionization modes using dynamic scheduled multiple reaction monitoring (MRM). The chromatographic separation employed two mobile phases: solvent A consisting of H2O/acetonitrile/isopropanol (50:30:20, v/v/v) containing 10 mM ammonium formate, and solvent B consisting of H2O/acetonitrile/isopropanol (1:9:90, v/v/v) also containing 10 mM ammonium formate. The gradient started at 10% solvent B with a flow rate of 0.4 mL/min, followed by sequential increases to 45%, 53%, 65%, 89%, 92%, and finally 100% B. Solvent B was maintained at 100% for 2.3 min before re-equilibration to initial conditions. The complete chromatographic run time was 16.5 min, including equilibration steps and transient flow rate adjustments up to 0.6 mL/min.

Lipid identification was based on specific precursor/product ion transitions together with retention time matching, as previously described (68). Peak integration was performed using Agilent MassHunter Quantitative Analysis software version 10.1. Relative quantification of lipid species was achieved by normalizing the peak area of each analyte to that of its corresponding internal standard (Supplementary Table 3).

#### 8.1. Lipid Ontology Enrichment Analysis

LION was used to identify altered lipid functions and organelle associations (https://www.lipidontology.com/ accessed 15 July 2026).

#### 8.2. CE saturation analysis

To investigate differences in saturation of CE between groups, the sum of the concentration of the CE with identical numbers of double bonds was calculated. These summed values were normalized over total lipid concentration. Afterward, the fold change from each sample was calculated over the mean of the control samples.

### 9. RNA Study

#### 9.1. RNA extraction

The RNA was extracted from the snap frozen frontal cortex from one hemisphere dissected as described in Sect. “Neurotransmitters analyses” and Sect. “Immunostainings” respectively, of *Tardbp*^+*/*+^ and *Tardbp^M323K/M323K^* male mice at 9 months of age (*n* = 4 per group). The tissue was homogenized with mechanical disaggregation in tubes with ceramic beads (Prescells) in 400 µl of Trizol. The RNA was extracted in Trizol and using columns following the protocol from the RNeasy Lipid Tissue Mini Kit (Qiagen) and removed potential genomic DNA contamination with DNAsa treatment in the columns (Qiagen), following the manufacturer’s instructions. The quantity and quality of the RNA were measured before use with a nanodrop and a qubit device (Thermofisher).

#### 9.2 Quantitative PCR analysis

Using the extracted RNA, cDNA synthesis was performed using the High-Capacity cDNA Reverse Transcriptase Kit (ThermoFisher Scientific) with 2 μg of total RNA. cDNA for qPCR reactions was used at a final concentration of 20 ng per well. Fast Sybr Green mastermix (ThermoFisher Scientific) was added, followed by the appropriate primer; final well volume of 10 μl. The primer of *Tardbp* to study the canonical isoforms were: Forward primer: 5’-AGGTTGCCCAGTCTCTTTGT-3’ and reverse primer:5’-GAACCTGATTCCCAAAGCCA-3’

#### 9.3 Short read RNA-sequencing and expression analysis

RNA was extracted from the frontal brain as described in Sect. “RNA extraction”, of wild-type and homozygous *Tardbp ^M323K/M323K^* male mice in the two diet regimes, at 9 months of age (n = 4 per group). Quality and quantity were assessed using the RNA Nano 6000 Assay Kit of the Bioanalyzer 2100 system (Agilent Technologies). Sequencing libraries were generated using an NEBNext® UltraTM RNA Library Prep Kit for Illumina® (NEB) following the manufacturer’s recommendations, and index codes were added to attribute sequences to each sample. The clustering of the index-coded samples was performed on a cBot Cluster Generation System using PE Cluster Kit cBot-HS (Illumina) according to the manufacturer’s instructions. After cluster generation, the library preparations were sequenced on an Illumina platform and paired-end reads were generated. The total number of reads was million per sample. Library preparation and RNA-seq was carried out by Novogene.

Quality control of FastaQ files was performed using FastQC (https://www.bioinformatics.babraham.ac.uk/projects). Low-quality reads (Phred quality score < 30) and reads too short (length < 15 pb) were removed using Fastp (69). The alignment to the genome (mm39 mouse reference genome) was achieved using HISAT2 for expression analysis (70).

The expression quantification of genes was carried out using FeatureCounts (71). Only uniquely mapped reads were used for the analysis of differential gene expression quantification with DESeq2 (72). Raw p-values were adjusted by the Benjamini–Hochberg false discovery rate (FDR) method and the adjusted p-values less than 0.05 were considered statistically significant. The pipeline is freely available in github.

Genes with an adjusted P-value <0.05 found by DESeq2 were assigned as differentially expressed. To identify which biological processes were altered from the RNA-seq study, ORA was used (73). ORA is a statistical method that determines whether genes from pre-defined sets, those belonging to a specific GO term, are present more than would be expected (overrepresented) in a subset of data. The ORA was performed using the clusterProfiler package (v3.16.1) in R. As input, it receives all (both upregulated and downregulated) DEGs from DESeq2, obtained using the cut-off criteria for statistical significance adjusted P-value <0.05.

#### 9.4. Multi-omics integration analysis

Transcriptomic and lipidomic datasets were integrated using the mixOmics package (v6.26.0) in R (v4.3.3). The transcriptomic dataset included normalized expression values of lipid metabolism-related genes, whereas the lipidomic dataset consisted of the relative abundance of each lipid species (% total lipids).

Multi-omics integration was performed using DIABLO (*block.splsda*), with experimental groups (*Tardbp^M323K/M323K^* CD and *Tardbp^M323K/M323K^* HFD) as the response variable. The model was built using two latent components, retaining 30 variables from each data block per component, and a design matrix with an inter-block connectivity value of 0.5. Data were centered and scaled to unit variance before analysis.

Associations between the selected transcriptomic and lipidomic features were visualized using Circos plots, displaying only correlations with an absolute correlation coefficient ≥ 0.90.

#### 9.5. Differential splicing analysis

To do the differential splicing analysis, we realigned the RNA-seq read using STAR (https://github.com/frattalab/rna_seq_snakemake). Then, alternative splicing events were identified and quantified by rMATS-turbo 4.3.0 (74) using gene transcript annotations from gencode mus musculus version mm39 rMATS was downloaded from the open-source platform SourceForge (https://rnaseq-mats.sourceforge.io/, accessed on 10 July 2025). Including novelSS events. The cut-off criteria for statistical significance were adjusted P-value <0.05 and |IncLevelDifference| > 5%.

Venn diagram was used in splicing and transcriptomic analysis to determine which genes or genes with splicing events were common between two or more lists of genes (75).

### 10. Statistical analysis

Statistical analyses of patients’ data were performed using Jamovi. Because neuropsychological variables were not normally distributed, non-parametric tests were used. Comparisons between two independent groups were performed using the Mann-Whitney’s U test. Associations between continuous variables were assessed using Spearman’s rank correlation coefficient (ρ). When appropriate, partial Spearman’s correlations were performed to adjust for confounding variables.

The statistical analyses for the mouse study were conducted using GraphPad Prism and RStudio. Two groups were compared with a single time point using Student’s t-test. Body weights between genotypes, across multiple time-points, were compared using 2-way ANOVA repeated measures/mixed models and Bonferroni correction of multiple testing. Two groups were compared across multiple time points using 2-way ANOVA with Sídak’s multiple comparisons post hoc test. Three or more groups were compared at a single time point using one-way ANOVA with Dunnett’s post hoc test. Statistical analysis of qRT-PCR data was performed on ΔCT values. Statistical significance was defined as p ≤ 0.05 for analysis of phenotyping and molecular biology data. Please see Figure legends for sample size - n numbers; n numbers refer to biological samples (i.e. number of animals used in animal experiments). Statistical detail for each experiment can be found in the Figure legends. Where indicated, * = p ≤ 0.05, ** = p ≤ 0.01, *** = p ≤ 0.001, **** = p ≤ 0.0001. Statistical analysis and Figures of Transcriptomic data were performed in RStudio using ggplot2, dplyr, DEGreport, reshape, ggrepel, RColorBrewer, pheatmap, ClusterProfiler, enrichplot and ggupset. All graphs were generated using GraphPad Prism 9. Venn diagrams were created using Venny2.1. BioRender.com was used to create original diagrams / figures.

## DECLARATIONS

### Ethics approval and consent to participate

The clinical study was approved by the Institutional Review Board (IRB) of Hospital Clínico San Carlos (approval no. 19/524-E_BS). Written informed consent was obtained from all individual participants included in the study.

The mouse study was approved by the local ethical committees of animal care and use of the Hospital Clínico San Carlos (Proex 222.7/24 and 258.4/20) in accordance with the European and Spanish regulations (2015/39/EU and RD 53/2013).

### Clinical trial number

Not applicable

### Availability of data and materials

Data are provided within the manuscript or Supplementary information files. Sequence data that support the findings of this study are deposited in GEO, openly accessible: GSE294236 and GSE344447.

### Authors’ contributions

Conceptualization: JMGC, IGT, JILC and SC. Formal Analysis: JMGC, IGT, JLC, ALS, ZA, JRN, PB, AF, MGM. Funding Acquisition: TC, PB and SC. Investigation: JMGC, IGT, LFH, LCFB, ZA, IJC, JJR, MGM, PB, RFR, MCF and AMA. Methodology: JMGC, IGT, ZA, PB, MPO, SC. Project administration: SC. Resources: PB, SC. Supervision: EF, TC, PB, MPO, JMGA, MGM and SC. Visualization: JMGC, IGT, PB, SC. Writing—original draft: JMGC, IGT, JILC and SC. Writing—review and editing: ALL.

## Acknowledgements

We thank all the patients who participated in this study for their time, dedication, and valuable contribution. We thank the nursing staff and technical personnel for their valuable support in the collection and processing of patient samples. We thank the animal care staff at the Hospital Clínico San Carlos (Madrid), Patricia Quesada and Cruz Bovada. We also thank the animal care staff at the Mary Lyon Centre at Harwell (UKRI) (Oxfordshire, UK). We thank the expert advice of Estela Area and for facilitating the lipidomic analysis.

## List of abbreviations

ACE-III: Addenbrooke’s Cognitive Examination III
AD: Alzheimer’s disease
ALS: Amyotrophic Lateral Sclerosis
BMI: body mass index
bvFTD: behavioural variant frontotemporal dementia
CD: Control Diet
CDR: Clinical Dementia Rating
CE: cholesterol esters
DEGs: differentially expressed genes
DG: diglycerides
ECAS: Edinburgh Cognitive and Behavioural
ALS: Screen
EE: energy expenditure
FAQ: Functional Activities Questionnaire
FTD: Frontotemporal Dementia
Hex1Cer: Hexosylceramide
HFD: High-fat diet
HOM: homozygous
IPGTT: intraperitoneal glucose tolerance tests
iWAT: inguinal white adipose tissue
LATE: limbic-predominant age related
TDP-43: encephalopathy
LPC: lysophosphocholines
NACC: National Alzheimer’s Coordinating Center
PC: phosphocholines
PUFAs: polyunsaturated fatty acids
RER: respiratory exchange ratio
SM: sphingomyelin
TG: triglycerides
WT: wild-type

