## Supplementary Figure 1 for "Loss of metabolic resilience underlies maladaptive responses to lipid overload in TDP-43 proteinopathy"

**Supplementary Material**

**Supplementary Table 1.** Partial correlations between serum metabolic biomarkers and cognitive performance assessed with ACE-III and ECAS in patients with bvFTD. Values are expressed as partial Spearman's correlation coefficients (ρ), with p-values shown in parentheses, adjusted for age and years of education.

| **Scores** | **Total cholesterol** | **HDL** | **LDL** | **Triglycerides** | **Prealbumin** | **HbA1c%** | **HOMA-IR** |
| --- | --- | --- | --- | --- | --- | --- | --- |
| **ACE-III (scaled)** | | | | | | | |
| **Total** | **-0.422 (0.028)** | 0.042 (0.840) | ***-0.350 (0.079)*** | ***-0.346***  ***(0.077)*** | ***-0.373 (0.088)*** | -0.063 (0.792) | 0.251 (0.216) |
| **Attention** | -0.268  (0.177) | 0.060 (0.771) | -0.232 (0.254) | **-0.400  (0.039)** | **-0.444 (0.038)** | **-0.452 (0.045)** | -0.010 (0.960) |
| **Memory** | -0.252 (0.204) | -0.087 (0.673) | -0.101 (0.625) | -0.092 (0.646) | -0.286 (0.197) | 0.296 (0.205) | 0.152 (0.458) |
| **Fluency** | **-0.492 (0.009)** | 0.090 (0.661) | **-0.483 (0.012**) | ***-0.327 (0.096)*** | ***-0.381 (0.080)*** | -0.197 (0.406) | 0.261 (0.198) |
| **Language** | -0.004 (0.982) | 0.228 (0.264) | 0.115 (0.577) | ***-0.355***  ***(0.069)*** | ***-0.414 (0.056)*** | -0.300 (0.198) | -0.048 (0.817) |
| **Visuospatial** | -0.105 (0.602) | 0.206 (0.312) | -0.152 (0.460) | -0.019 (0.925) | -0.311 (0.160) | -0.028 (0.905) | 0.346 (0.083) |
| **ECAS (age- and education-adjusted)** | | | | | | | |
| **Total** | -0.200 (0.425) | 0.143 (0.583) | -0.069 (0.791) | **-0.478 (0.045)** | -0.202 (0.508) | -0.072 (0.823) | -0.114 (0.663) |
| **Language** | -0.160 (0.526) | 0.193 (0.458) | 0.043 (0.869) | **-0.492 (0.038)** | -0.149 (0.628) | -0.156 (0.629) | -0.234 (0.366) |
| **Fluency** | -0.265 (0.289) | -0.440 (0.077) | -0.105 (0.688) | -0.219 (0.382) | 0.016 (0.958) | 0.272 (0.393) | 0.223 (0.390) |
| **Executive** | -0.144 (0.569) | 0.046 (0.860) | -0.131 (0.617) | -0.440 (0.068) | -0.441 (0.132) | -0.517 (0.085) | -0.054 (0.837) |
| **Memory** | -0.063 (0.805) | 0.311 (0.224) | 0.044 (0.867) | -0.293 (0.238) | -0.150 (0.625) | 0.164 (0.610) | -0.133 (0.610) |
| **Visuospatial** | 0.179 (0.477) | 0.458 (0.065) | 0.412 (0.101) | ***-0.421 (0.082)*** | -0.059 (0.848) | -0.187 (0.561) | -0.374 (0.139) |

**Supplementary Table 2.** Antibody list.

| **PRODUCT** | **SUPPLIER** | **REFERENCE** |
| --- | --- | --- |
| **HOESCHT** | ThermoFisher Scientific | H21492 |
| **GFAP Antibody** | Cell Signaling | 36565 |
| **Iba1 Antibody** | Abcam | ab5076 |
| **Plin2 Antibody** | Bio-Techne (Novus Biologicals) | NB110-40877 |
| **Alexa 488 rabbit** | Invitrogen | A32731 |
| **Alexa 488 mouse** | Invitrogen | A32723 |
| **Alexa 594 rabbit** | Invitrogen | A11037 |
| **Alexa 594 mouse** | Invitrogen | A11032 |
| **Donkey Anti-Goat 555** | Abcam | Ab150134 |

**Supplementary Table 3.** Class-representative internal lipid standards.

| **ISTD** | | **Reference** | **Commercial supplier** |
| --- | --- | --- | --- |
| AcylCarnitine(18:1) d9 | AcylCarnitine(18:1) | 870321P | Avanti polar lipids |
| CE | CE mixture | 330822L | Avanti polar lipids |
| Cer (d18:1/17:0) | Ceramide (d18:1/17:0) | 860517P | Avanti polar lipids |
| CHO d7 | Cholesterol d7 | 700041P | Avanti polar lipids |
| DG (16:1) d5 | Diacylglycerol (16:1) d5 | 800853C | Avanti polar lipids |
| dhCer (d18:0-d7/13:0) d7 | Dihydroceramide (d18:0-d7/13:0) d7 | 330726P | Avanti polar lipids |
| FA 22:6 d5 | Fatty acid (22:6= d5 | 733326 | Avanti polar lipids |
| Hex1Cer (d18:1-d7/15:0) | Glucosyl(β) Ceramide-d7 (d18:1-d7/15:0) | 330729P | Avanti polar lipids |
| Hex2Cer (d18:1-d7/15:0) | Lactosyl(β) Ceramide-d7 (d18:1-d7/15:0) | 330727P | Avanti polar lipids |
| LPC(13:0) | Lyso PC (13:0) | 855476P | Avanti polar lipids |
| LPE (18:1) d7 | Lyso PE (18:1) d7 | 791644C | Avanti polar lipids |
| PC(13:0) | Phosphatidylcholine (13:0) | 850340P | Avanti polar lipids |
| PE(17:0) | Phosphatidylethanolamine (17:0) | 830756P | Avanti polar lipids |
| PG(17:0) | Phosphatidylglycerol (17:0) | 830456P | Avanti polar lipids |
| SM (18:1/18:1) d9 | SM (d18:1/18:1) d9 | 860740L | Avanti polar lipids |
| TG (51:0) | Triglyceride (51:0) | T2151 | Sigma Aldrich |

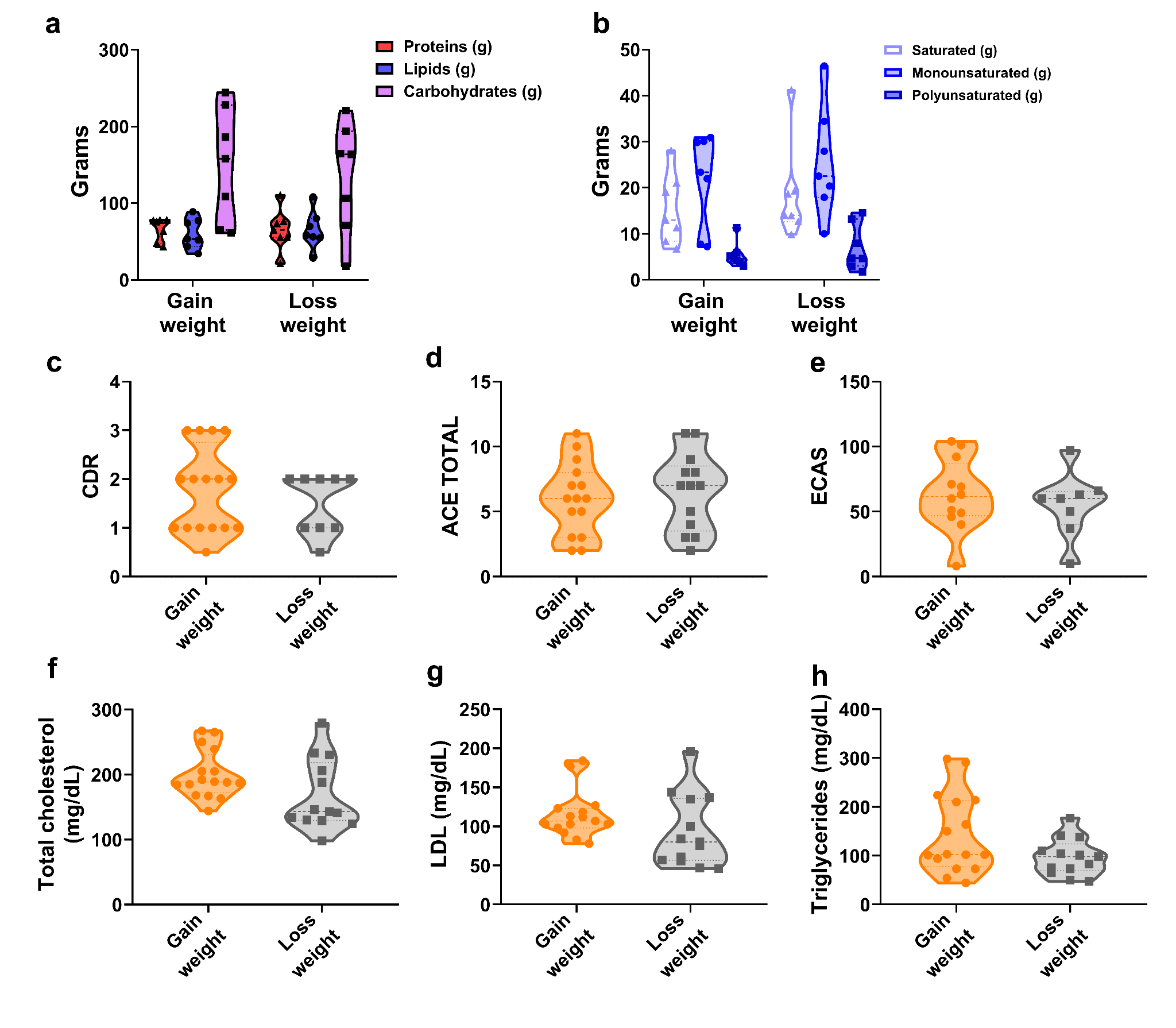

**Supplementary Figure 1.** ***a.*** *Quantitative composition of daily macronutrient intake (grams of protein, lipids, and carbohydrates).* ***b.*** *Lipid profile breakdown, demonstrating the relative distribution of saturated, monounsaturated, and polyunsaturated fatty acids.* ***c.*** *Baseline assessment of cognitive and functional status using the Clinical Dementia Rating (CDR) scale.* ***d****. Global cognitive performance assessed by the Addenbrooke's Cognitive Examination III (ACE-III) total score.* ***e****. Total score on the Edinburgh Cognitive and Behavioural ALS Screen (ECAS).* ***f, g, h.*** *Serum metabolic markers showing total cholesterol (f), LDL (g) and triglyceride levels (h) measured in mg/dL.*

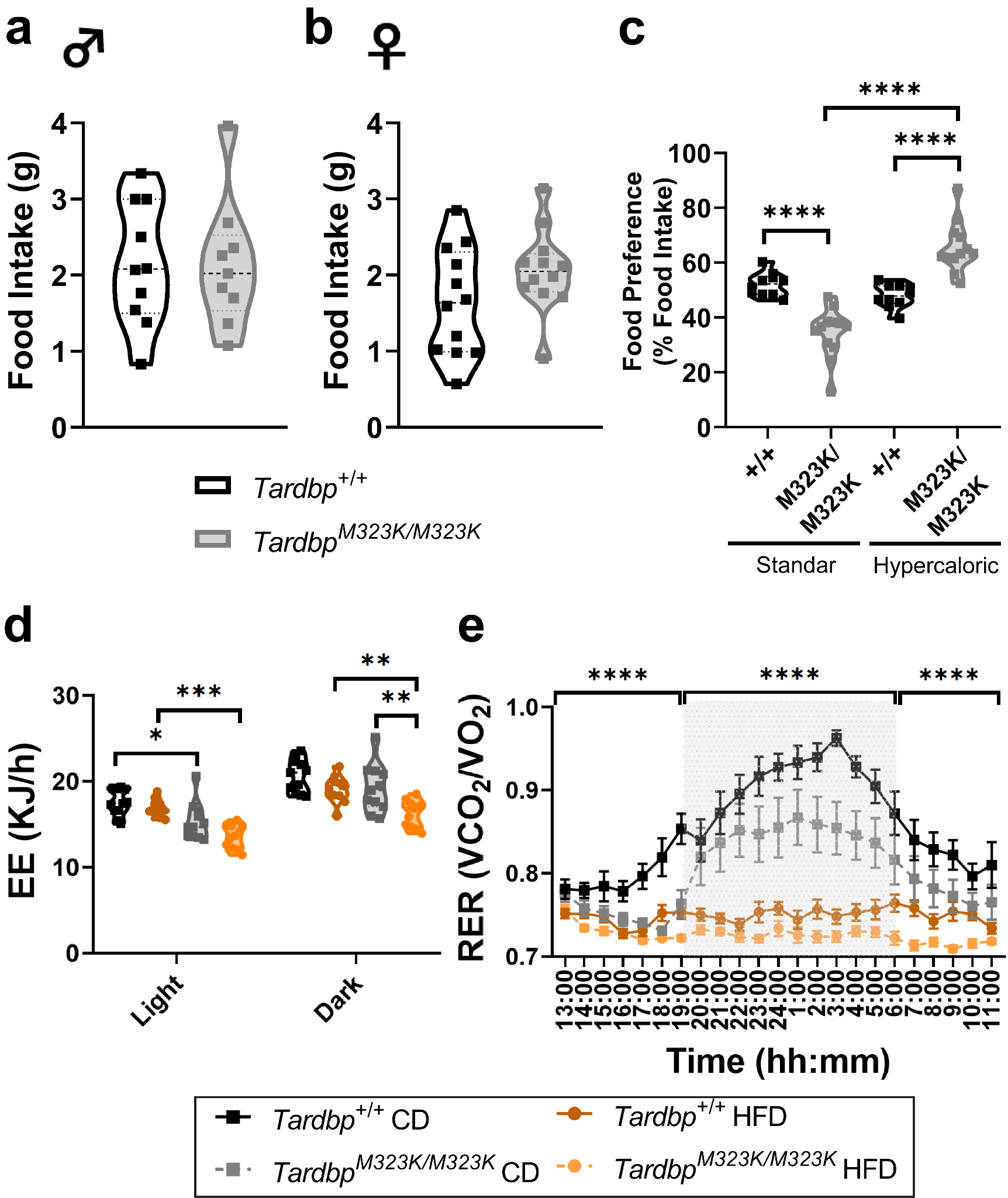

***Supplementary Figure 2. Metabolic and Cognitive impact of High Fat Diet in a TDP-43 proteinopathy model. a.*** *Food intake in 9-month-old males, n: Tardbp^+/+^ = 10, Tardbp^M323K/M323K^ = 9* ***b.*** *Food intake in 9 month-old females, n: Tardbp^+/+^ = 12, Tardbp^M323K/M323K^ = 12.* ***c.*** *Food preference in 9 month old females, n: Tardbp^+/+^ = 12, Tardbp^M323K/M323K^ = 12.* ***d****. Indirect calorimetry, energy expenditure measurements; n: Tardbp^+/+^ CD = 11, Tardbp^M323K/M323K^ CD = 11, Tardbp^+/+^ HFD = 11, Tardbp^M323K/M323K^ HFD = 11.* ***e****. Indirect calorimetry, respiratory exchange ratio (RER); n: Tardbp^+/+^ CD = 11, Tardbp^M323K/M323K^ CD = 11, Tardbp^+/+^ HFD = 11, Tardbp^M323K/M323K^ HFD = 11.*

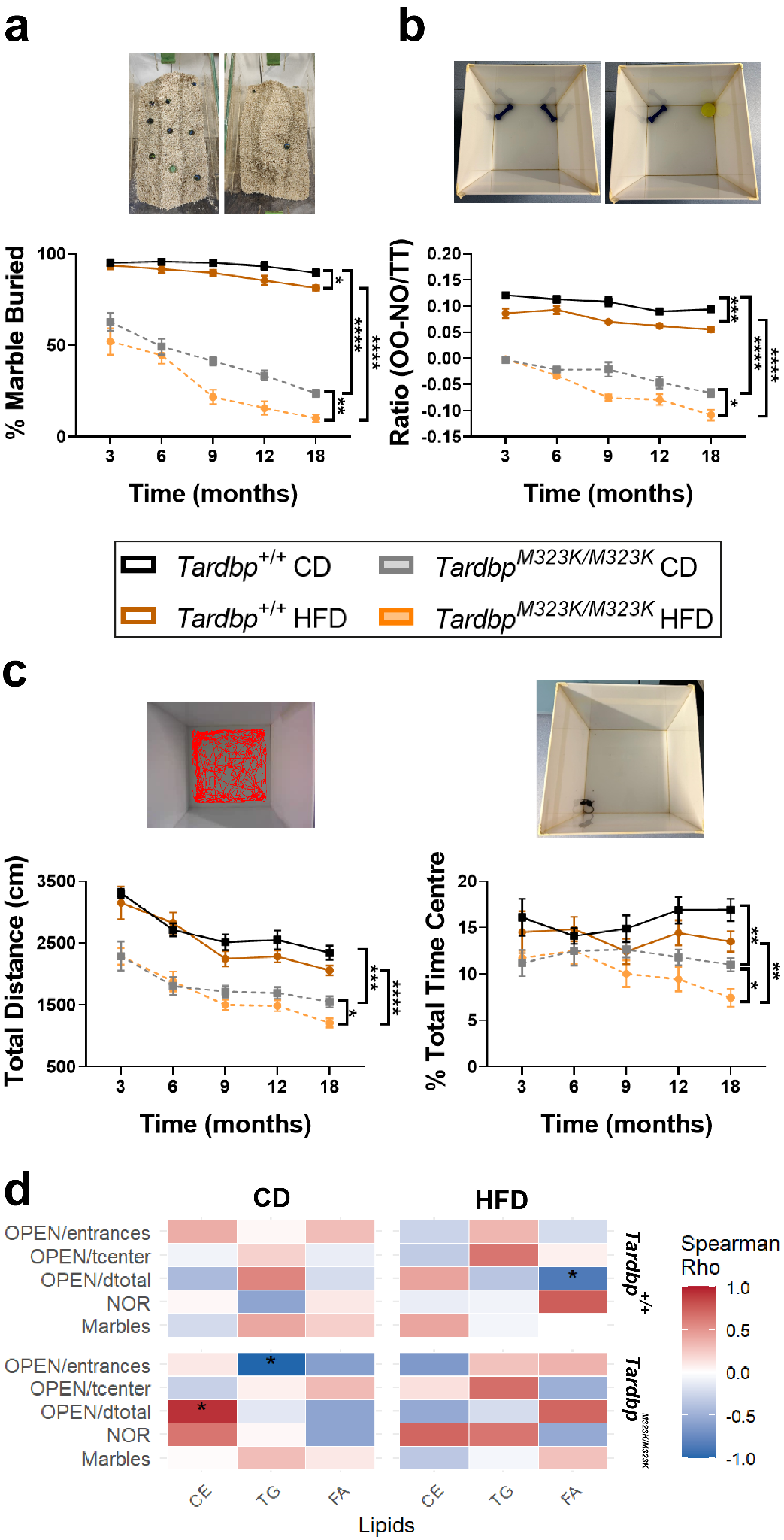

***Supplementary Figure 3.*** ***Lifelong high fat diet effects on the cognitive induced by the Tardbp ^M323K/M323K^ mutation in females.*** ***a****. Marble burying test in females up to 18 months of age.* ***b****. Novel Object Recognition (NOR) test in males up to 18 months of age.* ***c****. Open field test in males up to 18 months of age, showing total distance traveled and time spent in the center.* ***d****. Correlation between lipid levels and different behavioural tests in males. (a-c: n: Tardbp^+/+^ CD = 9, Tardbp^M323K/M323K^ CD = 7, Tardbp^+/+^ HFD = 8, Tardbp^M323K/M323K^ HFD = 11 in females).*

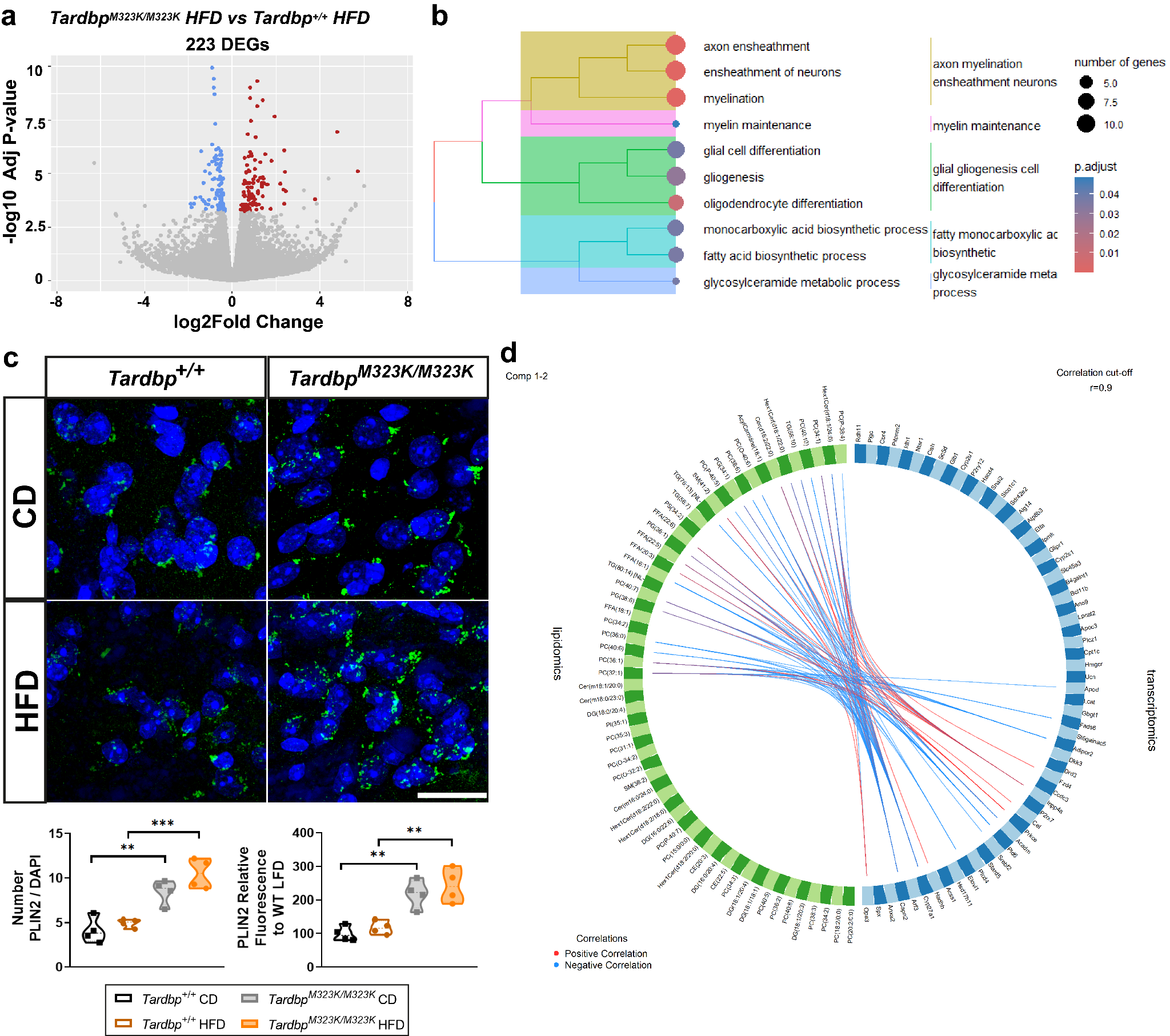

***Supplementary Figure 4.*** ***a****. Volcano plot of the 223 DEGs between the Tardbp^M323K/M323K^ HFD vs Tardbp^M323K/M323K^ CD comparison.* ***b.*** *Over-representation analysis (ORA) of deregulated genes.* ***c****. Representative image and quantification of PLIN2 immunostaining (PLIN2 in green and Hoetsch in blue) in the frontal cortex of 9-month-old males; n: 4 per group. Scale bar = 20 µm.* ***d****. Mixomics integration between lipidomics and transcriptomics.*
